# The INO80 chromatin remodeling complex regulates histone H2A.Z mobility and the G1-S transition in oligodendrocyte precursors

**DOI:** 10.1101/2024.01.09.574847

**Authors:** Jordan L Wright, Yi Jiang, Stuart Nayar, Vinothini Rajeeve, Pedro R Cutillas, Huiliang Li, William D Richardson

## Abstract

Chromatin remodelling complexes (CRCs) participate in oligodendrocyte (OL) differentiation, survival and maintenance. We asked whether CRCs also control proliferation of OL precursors (OPs) – focusing on the INO80 complex, which is known to regulate proliferation of a variety of other cell types during development and disease. CRISPR/Cas9-mediated inactivation of *Ino80* in vitro, or Cre-mediated deletion in vivo, slowed the OP cell cycle substantially by prolonging G1, without inducing OL differentiation. RNAseq analysis revealed that E2F target genes were dysregulated in OPs from INO80-deficient mice, but correlated RNAseq and ATAC-seq uncovered no general correlation beween gene expression and altered nucleosome positioning at transcription start sites. Fluorescence photobleaching experiments in cultured OPs demonstrated that histone H2A.Z mobility increased following loss of INO80, suggesting that INO80 regulates the cell cycle machinery in OPs through H2A.Z/ H2A exchange. We also present evidence that INO80 associates with OLIG2, a master regulator of OL development.

## INTRODUCTION

Oligodendrocytes (OLs) synthesize the myelin sheaths around central nervous system (CNS) axons, increasing the speed of action potentials while reducing the associated energy demand on axons (Hartline and Colman, 2007). Myelinating OLs are also believed to transfer substrates for energy production (e.g. lactate) into the axons that they ensheath (Saab et al., 2013) OLs are generated from OL precursor cells (OPs) that originate in the ventricular zone (VZ) of the embryonic brain and spinal cord, then proliferate and migrate through the developing CNS before differentiating into myelin-forming OLs. A substantial population of OPs persists in the grey and white matter of the postnatal CNS, continuing to self-renew and differentiate into new myelinating OLs throughout young adult life (Richardson et al., 2011; Nishiyama et al., 2021).

The decision of an OP to divide, or to exit the cell cycle and differentiate into an OL, requires the transcriptional activation or repression of different sets of genes. This depends on key transcription factors (TFs) such as OLIG2, SOX10 and MYRF, which act together or sequentially to orchestrate different stages of OL development from embryonic neural stem cell to mature myelinating OL (Elbaz and Popko, 2019). Of these, OLIG2 stands out because it is expressed and required at every stage of OL development – unlike MYRF, for example, which is expressed only in differentiating and mature OLs (Elbaz and Popko, 2019; Sock and Wegner, 2021). It is possible that OLIG2’s special status as a master regulator results from stage-specific interactions with chromatin remodeling complexes (CRCs), with which it cooperates to direct site- and stage-specific remodeling of nucleosomes along the genome (Yu et al., 2013; He et al., 2016; Marie et al., 2018; Zhao et al., 2018; Elsesser et al., 2019; Parras et al., 2020).

CRCs are large, multi-subunit assemblies, each of which contains a single ATPase motor protein that facilitates nucleosome sliding, eviction, recruitment or histone subunit exchange, all in an ATP-dependent manner. There are at least four CRCs in mammals, classified on the basis of their intrinsic ATPases: 1) the BAF/ SWI/ SNF complex (BRG1 or BRM ATPase), 2) the NuRD/ CHD complex (one of CHD3/4/5 ATPases), 3) the ISWI complex (SNF2L or SNF2H ATPase) and 4) the INO80 complex (one of INO80/ SWR family ATPases). Germline deletion of any of the intrinsic ATPases results in early embryonic lethality, indicating essential roles for CRCs during development (Jin et al., 2015; Zhou et al., 2016a; Zhou et al., 2016b; Alvarez-Saavedra et al., 2019). CRC subunit genes continue to be expressed throughout life in a broad range of tissues and cell types.

Several labs have explored the functions of CRCs in the OL lineage. OL lineage-specific deletion of BRG1, a BAF complex ATPase, inhibits OL differentiation and myelination to a greater or lesser extent (Yu et al., 2013; Bischof et al., 2015), possibly depending on the choice of CRE driver and hence the precise stage of OL lineage development at which deletion occurs (Matsumoto et al., 2016). Consistent with these studies, conditional knockout (cKO) of CHD8 (Zhao et al., 2018) or CHD7 (He et al., 2016; Marie et al., 2018), which act respectively upstream or downstream of BRG1, inhibit OP proliferation and survival – as does cKO of BAF complex scaffold proteins BAF155 and BAF170, potentially inactivating both BRG1- and BRM-containing complexes (Abbas et al., 2021). Other CRCs have been implicated in the control of OL lineage progression; for example, cKO of both HDAC1 and HDAC2, histone deacetylases associated with the NuRD complex (Ye et al., 2009), inhibits OL differentiation, while cKO of EP400, an ATPase of the INO80/ SWR family, inhibits OL survival and myelination (Elsesser et al., 2019). Together, these studies suggest that different CRCs perform essential roles at different stages of the OL lineage.

With the exception of CHD8 (Zhao et al., 2018), those CRCs that have been explored in the context of OL development have been found not to impact OP proliferation. However, the INO80 complex is known to play an important role in proliferation of other cell types (Shimada et al., 2008; Qiu et al., 2016; Kokavec et al., 2017; Knezevic et al., 2018; Rhee et al., 2018; Alvarez-Saavedra et al., 2019), including promoting tumorigenicity in various cancers (Jin et al., 2015; Zhou et al., 2016a; Zhou et al., 2016b). INO80 is recruited to E2F target genes (encoding proteins required for the G1-S transition) and deletion of proteins of the INO80 complex results in reduced and dysregulated E2F target gene expression (Knezevic et al., 2018). Histone variants are also known to be critical at specific phases of the cell cycle. For example, deletion of histone variant *H2A.Z* results in an elongated cell cycle with delayed S-phase enry linked to delayed expression of G1- and S-phase cyclin genes (Dhillon et al., 2006). The INO80 complex is known to facilitate H2A.Z– H2A exchange in nucleosomes (Papamichos-Chronakis et al., 2011; Alatwi and Downs, 2015; Brahma et al., 2017), therefore we hypothesized that this function of INO80 might play an important role in controlling the OP cell cycle. We disabled the INO80 complex specifically in the OL lineage through conditional deletion of the INO80 ATPase gene using CRISPR/Cas9-mediated gene disruption in vitro and Cre-lox recombination in vivo. This identified the INO80 complex as a key cell cycle regulator in proliferating OPs, seemingly with no direct role in promoting OL differentiation. Fluorescence photobleaching experiments in cultured OPs demonstrated that *Ino80* deletion resulted in increased mobility of histone H2A.Z in the nucleus, implying that the INO80 complex is important for H2A.Z-dependent nucleosome remodeling in cycling OPs. In addition, *Ino80* deletion in OPs resulted in altered expression of E2F target genes, which are important regulators of the G1 to S-phase transition.

## RESULTS

### Dynamic INO80 expression as OPs differentiate into OLs

We immunolabelled for INO80 in postnatal day 7 (P7) mouse brain and found that INO80 is detectable in all cell nuclei observed in our brain sections, including nuclei of OLIG2^+^ OL lineage cells in the grey and white matter (Fig. 1A). We confirmed that OL lineage cells express INO80 by Western blot of protein extracts from immuno-purified OPs from P7 mice expanded *in vitro* in the presence of platelet-derived growth factor (PDGF)-AA or induced to differentiate into OLs in the presence of thyroid hormone (TH) (Fig. 1B-C). INO80 protein level was highest in proliferating OPs but decreased within a day of inducing cell cycle exit and differentiation, remaining at a constant low level thereafter (Fig. 1C).

**Figure 1.**
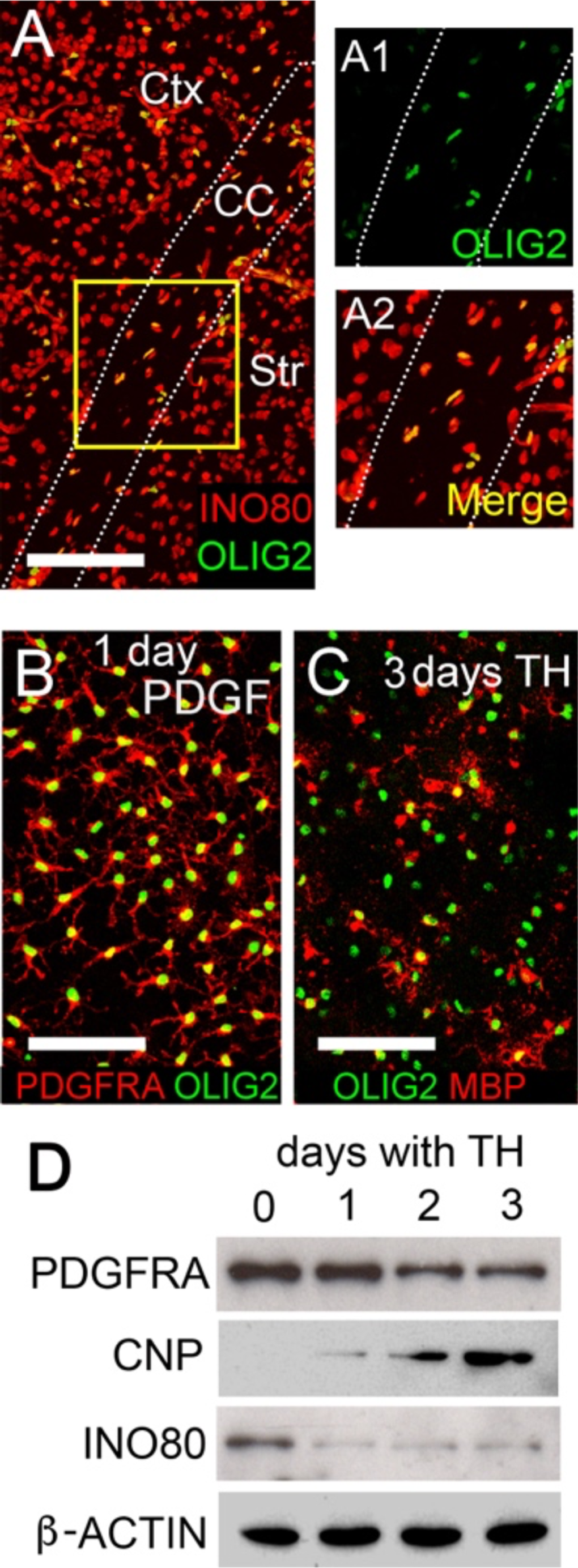
Dynamic regulation of *Ino80* during OL differentiation. (**A**) Coronal section of P0 mouse forebrain immunolabelled for INO80 and OLIG2. Higher-magnification images of the indicated area are shown in **A1**, **A2**. INO80 is co-expressed with OLIG2 in OL lineage cells in the grey and white matter. Ctx, cortex; Str, striatum; CC, corpus callosum. (**B, C**) Immuno-purified OPs cultured for 24 hours in the presence of PDGF and absence of thyroid hormone (TH) (**B**), or three days in the presence of TH (**C**), immunolabelled as shown. MBP-positive differentiated OLs are present after 3 days exposure to TH. (**D**) Western blots of whole-cell protein extracts of OPs cultured with TH for 1-3 days and probed with the indicated antibodies. An OP marker (PDGFRA) and differentiated OL marker (2’,3’-cyclic nucleotide 3’-phosphodiesterase, CNP) were included to assess the differentiation state of the cultures, along with a loading control (β-Actin). INO80 is rapidly down-regulated following cell cycle exit and initiation of OL differentiation. Scale bar: 100 µm (**A**), 50 µm (**B, C**).

### INO80 is required for OP proliferation in vitro

As INO80 is an essential ATPase subunit of the INO80 complex, *Ino80* gene disruption should block the ATP-dependent nucleosome remodelling function of this complex. Therefore, to ask whether the INO80 complex is functionally important during OL lineage development we used CRISPR-CAS9 to introduce inactivating mutations into the *Ino80* gene in cultured OPs. We tested two different guide RNAs (*Ino80.g1* and *Ino80.g2*), both of which resulted in disruption of the *Ino80* gene based on the surveyor assay (Fig. 2A), as well as markedly reducing protein levels on Western blots when compared to *Cas9*-only controls (Fig. 2B). Using these guide RNAs we targeted *Ino80* in OP cultures and added 5-ethynyl-2-deoxyuridine (EdU) to the culture medium for 6 hours to quantify OP proliferation. *Ino80* targeted OP cultures contained a greatly reduced proportion of EdU^+^ cells relative to *Cas9*-only controls, (Fig. 2C,D). To assess the potential for OPs to differentiate into OLs in the absence of INO80 we induced OL differentiation by withdrawing PDGF-AA and adding TH to the culture medium for three days. We then immunolabelled the cultures for myelin basic protein (MBP), which marks differentiated OLs. There was no difference in the proportion of cells that expressed MBP in *Ino80*-depleted cultures compared to *Cas9*-only controls (Fig. 2E,F), suggesting that the probability of an OP differentiating to a post-mitotic OL is not dependent on INO80, despite the reduced division rate of OPs. This might be because loss of INO80 causes OPs to arrest at the G1/S boundary without committing to leave the cell cycle and differentiate.

**Figure 2.**
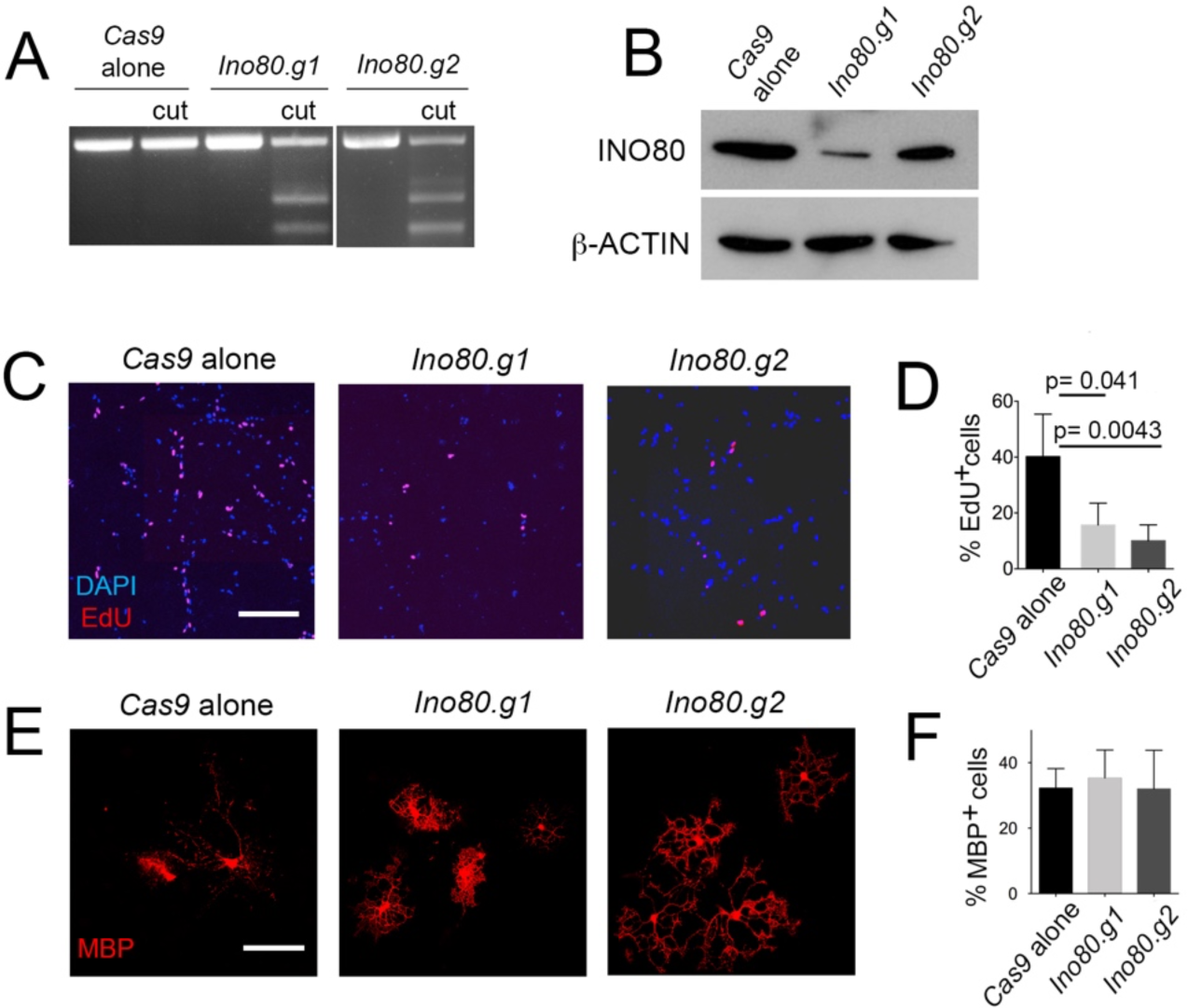
CRISPR/ Cas9-mediated disruption of *Ino80* in immuno-purified OPs. (**A, B**) Assessment of gene disruption by (**A**) “Surveyor assay” (see Methods) and (**B**) Western blot, after transfection of cultured immuno-purified OPs with *Cas9* + guide RNAs (gRNAs) targeting *Ino80*, and subsequent puromycin treatment. Two different gRNAs (*Ino80.g1* and *Ino80.g2*) were compared to a *Cas9*-only control (see Table 1 for gRNA sequences); *Ino80.g1* appeared more effective than *Ino80.g2*. (**C**) OPs transfected with Cas9 alone (control), *Cas9* + *Ino80.g1* or *Cas9* + *Ino80.g2* were treated for 6 h with EdU in culture medium containing PDGF, before fixing the cells and labelling for EdU and DAPI. *Ino80*-KOs had visibly fewer EdU-positive cell nuclei than controls. (**D**) Quantifying the fraction of OPs that incorporated EdU revealed a significant reduction in cell division rate in *Ino80*-KOs. (**E**) *Ino80*-KO OPs were cultured in the presence of TH for 3 days to induce OL differentiation, before immunolabelling for Myelin basic protein (MBP). (**F**) Quantifying the fraction of OPs that expressed MBP revealed no significant differences between *Cas9*-only and *Ino80*-KOs. Error bars represent mean ± s.e.m.. Statistical significance is based on 1-way ANOVA with post-hoc Dunnett’s multiple comparison test. Scale bars: 50 µm (**C**, **E**).

**Table 1.**
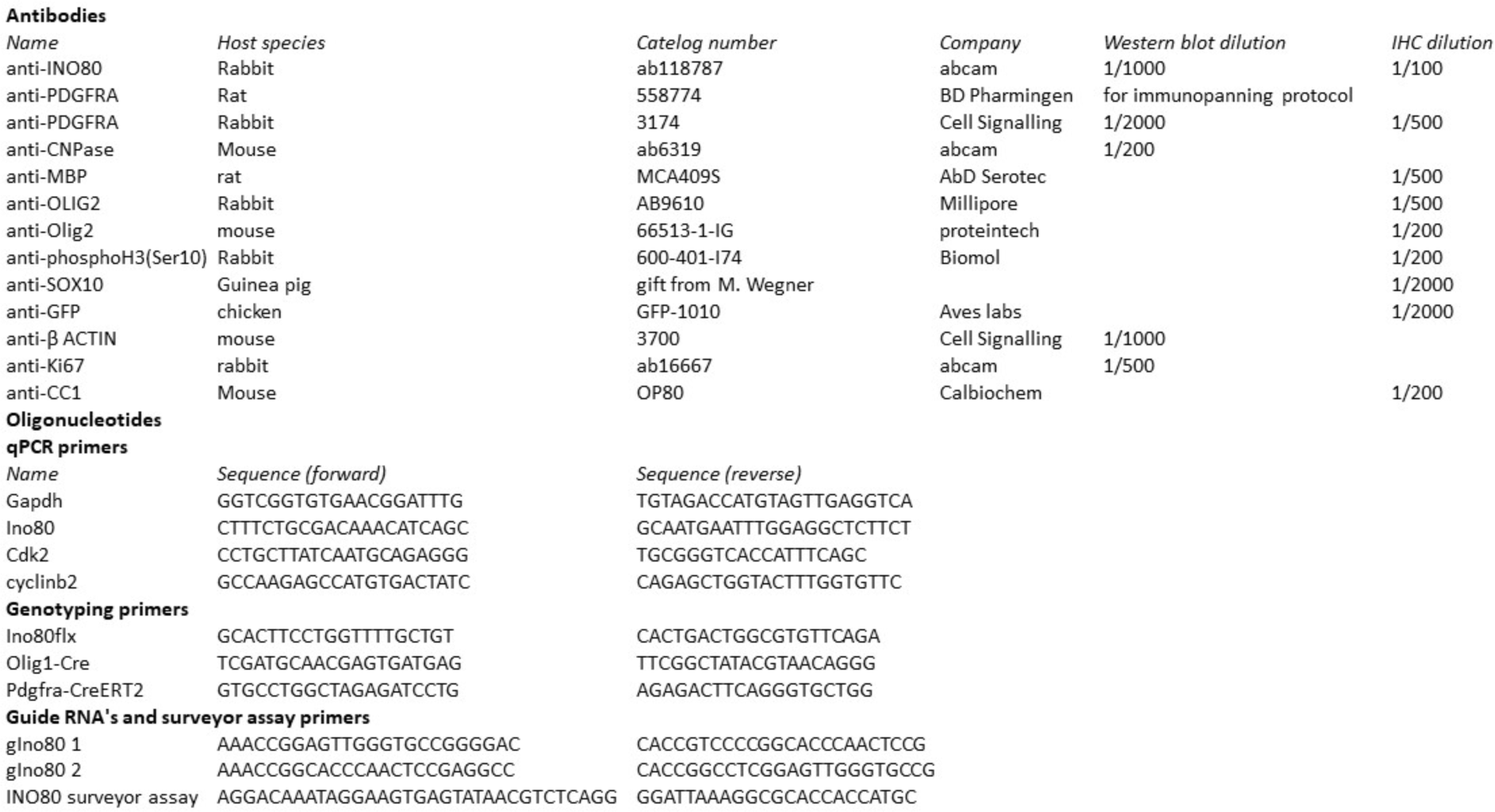
List of primary antibodies and DNA oligonucleotides.

### Conditional knockouts confirm that INO80 is required for OP proliferation in perinatal mice

To validate our in vitro results in animal models we ablated INO80 specifically in the OL lineage, by crossing *Olig1-Cre* mice with a “floxed” allele of *Ino80 (Ino80^flx^)* to generate *Olig1-Cre: Ino80^flx/flx^* offspring (referred to as *Olig1-Ino80*-cKOs) (Supplementary Fig. 1A). Controls were *Ino80^flx/flx^* without *Cre* and/or *Olig1-Cre: Ino80^flx/+^* heterozygotes. *Olig1-Cre: Rosa26-YFP* mice confirmed that CRE-mediated recombination was limited to OLIG2+ OL lineage cells in the brain (Supplementary Fig. 1B-C). CRE-recombination at the *Ino80* locus, assayed by quantitative PCR (qPCR) of reverse-transcribed RNA from cultured OPs, reduced *Ino80* mRNA levels to less than one-third of normal (Supplementary Fig. 1D). We examined OP development in the corpus callosum underlying the motor cortex at P0, P7 and P14, when the number-density of OPs is high. The density of OPs was significantly reduced in *Olig1-Ino80*-cKOs at P0 and P7 relative to controls (P0: 85 ± 5 OPs/mm^2^ in *Ino80*-cKOs vs 205 ± 20 OPs/mm^2^ in controls, p=6.7 x 10^−4^. P7: 177 ± 35 OPs/mm^2^ in *Ino80*-cKOs vs 290 ± 15 OPs/mm^2^ in controls, p=0.013. Means ± s.e.m). By P14, although there were still fewer OPs in *Olig1-Ino80*-cKOs, this was no longer significant (P14: 111 ± 18 OPs/mm^2^ in *Ino80*-cKOs vs 150 ± 19 OPs/mm^2^ in controls, p=0.21) (Fig. 3A,B,D). To determine whether the early reduction in the number of OPs reflected a proliferation deficit, as observed in cultured OPs, the fraction of OPs in vivo that was actively engaged in the cell cycle (i.e. not arrested in early G1) was assessed by co-immunolabeling for PDGFRA and Ki67. A smaller percentage of PDGFRA^+^ OPs was actively cycling (Ki67^+^) in *Olig1-Ino80*-cKOs compared to controls both at P0 and P7 (P0: 39% ± 8% Ki67^+^ OPs/mm^2^ in *Ino80*-cKOs vs 57% ± 6% in controls, p=0.006. P7: 31% ± 4% Ki67^+^ OPs/mm^2^ in *Ino80*-cKOs vs 42% ± 7% in controls, p=0.004. Means ± s.e.m) (Fig. 3C). At P14 there was also a reduction although this did not reach statistical significance (P14: 50% ± 11% Ki67^+^ OPs/mm^2^ in *Ino80*-cKOs vs 34% ± 3% in controls, p=0.066. Means ± s.e.m) (Fig. 3C). An INO80-dependent proliferation deficit in newborn mice was confirmed by administering a single injection of EdU and analyzing the mice 2 h later to determine the fraction of PDGFRA^+^ OPs in the corpus callosum that was in S-phase (EdU^+^) (labeling index, L.I.=0.16 ± 0.02 in *Ino80*-cKOs versus 0.33 ± 0.02 in controls, p<0.001) (Fig. 3E). This demonstrates that S-phase occupies a smaller proportion of the total cell cycle in *Olig1-Ino80*-cKOs than in controls — most likely because G1 is prolonged in the cKOs (see below).

**Figure 3.**
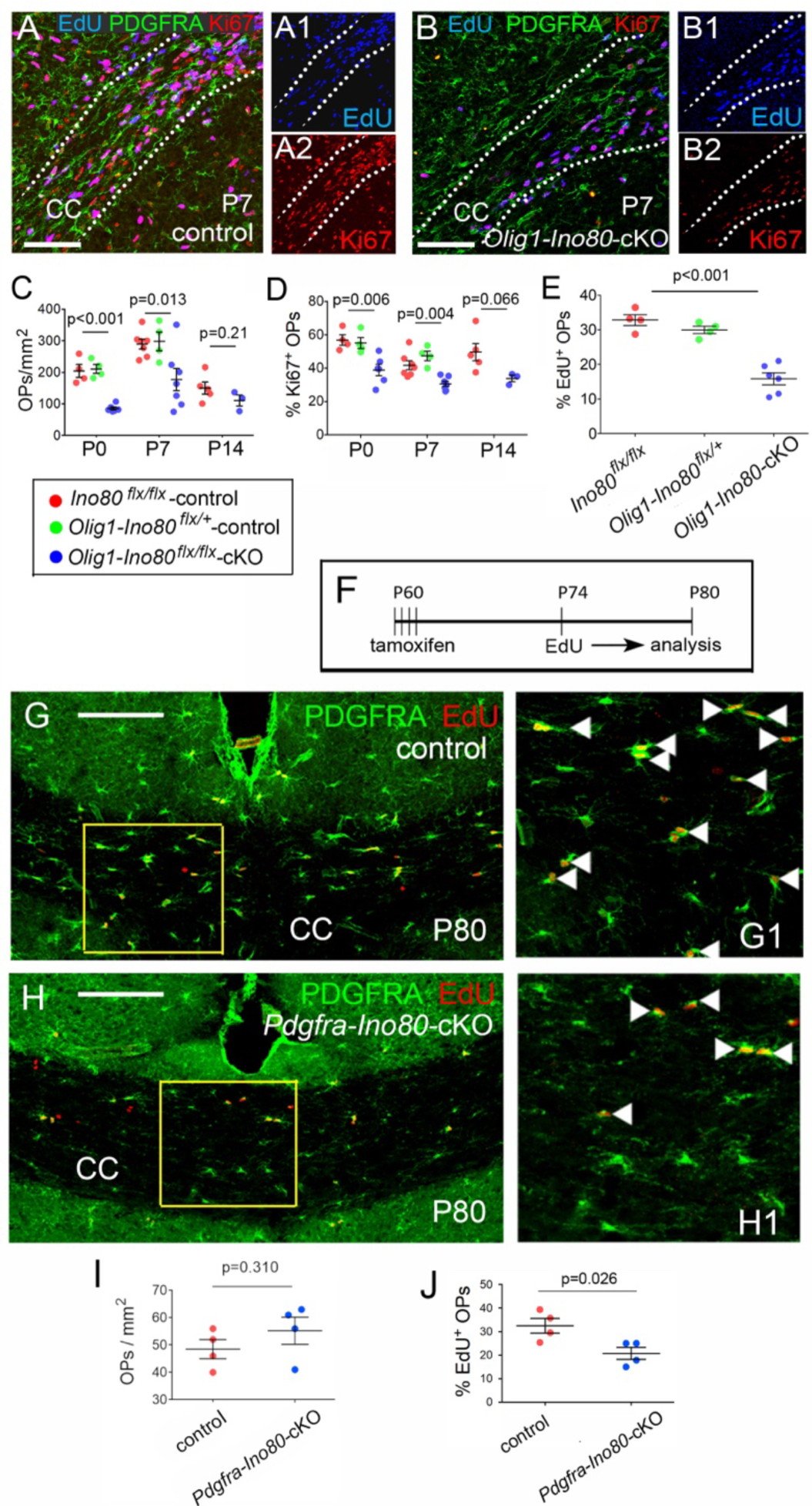
Reduced OP proliferation in perinatal and adult *Ino80*-cKO mice. (**A**) *Olig1-Cre*: *Ino80^flx/+^* or *Ino80^flx/flx^* mice (controls) and (**B**) *Olig1-Cre*: *Ino80^flx/flx^* (*Olig1-Ino80*-cKO) mice were each given a single injection of EdU at P0, P7 or P14. 4 h later they were killed and coronal forebrain sections labelled by EdU histochemistry combined with immunolabelling for Ki67 and PDGFRA (*Olig1-Cre: Ino80^flx/+^* controls are shown). Dotted lines delineate the corpus callosum (CC). Separate EdU (blue) and Ki67 (red) channels are shown in **A1**,**A2** and **B1**,**B2**. There were visibly fewer (EdU^+^, Ki67^+^, PDGFRA^+^) OPs in *Olig1-Ino80*-cKOs than in control littermates. (**C**) Counting total PDGFRA^+^ OPs in the CC revealed significantly fewer OPs in *Olig1-Ino80*-cKOs compared to littermate controls at P0 and P7 but not at P14. (**D**) Quantifying the fraction of PDGFRA^+^ OPs that was Ki67^+^ in the CC revealed significantly fewer Ki67^+^ (cycling) OPs in *Olig1-Ino80*-cKOs relative to littermate controls at P0 and P7 but not at P14. (**E**) The fraction of PDGFRA^+^ OPs that was EdU^+^ in the CC was also significantly smaller in *Olig1-Ino80*-cKOs compared to *Ino80^flx/flx^*control littermates at P7. (**F**) Adult (P60) *Ino80^flx/flx^* control mice and (**G**) *Pdgfra-Ino80*-cKO littermates were given tamoxifen by gavage (see **H** for protocol), then provided with EdU in their drinking water for one week P74-P80 before immunolabelling for PDGFRA along with EdU histochemistry. (**I**) Quantifying the fraction of PDGFRA^+^ OPs that was also EdU^+^ showed that the proportion of OPs that incorporated EdU was significantly reduced in *Pdgfra-Ino80*-cKOs relative to control littermates. Error bars represent mean ± s.e.m. p-values are based on 1-way ANOVA with post-hoc Dunnett’s multiple comparison test and Student’s t-test. Scale bars: 100 µm (**A**, **B**), 200 µm (**G**, **H**).

The OP cell cycle slows down steadily with age in mice, e.g. from ∼3 days in the corpus callosum at P21 to ∼10 days at P60, but all OPs remain capable of dividing at P60 because they can all incorporate EdU, given enough time (Young et al., 2013). We conditionally deleted *Ino80* in adult OPs (sparing differentiated OLs) by administering tamoxifen to *Pdgfra-CreER^T2^: Ino80^flx/flx^*mice (referred to as *Pdgfra-Ino80*-cKOs) at P60-64. We subsequently labeled dividing OPs by administering EdU in the drinking water for one week P74-P80, then immunolabelled forebrain sections for PDGFRA together with EdU histochemistry (Fig. 3F-H). We found that the EdU labeling index of PDGFRA^+^ OPs in the corpus callosum was reduced significantly in *Pdgfra-Ino80*-cKO adult mice compared to control littermates (L.I. = 0.21 ± 0.03 in *Pdgfra-Ino80*-cKOs vs 0.32 ± 0.06 in controls, p=0.026) (Fig. 3I), without significant reduction in the number-density of OPs (Fig. 3F,G).

### Ino80-cKO causes a transient developmental delay in OL production

We examined the production of OLs in *Olig1-Ino80*-cKO mice. We counted the number-density of mature (CC1^+^, OLIG2^+^) double-labeled OLs in the anterior corpus callosum at P7 (Fig. 4A). We found a reduced density of OLs in *Olig1-Ino80*-cKOs compared to controls (22 ± 3 OLs/mm^2^ in *Ino80*-cKOs vs 57 ± 4 in controls, p=0.007) (Fig. 4B). However, there was no difference in the proportion of all OL lineage cells (OLIG2^+^) that was (CC1^+^, OLIG2^+^) mature OLs (Fig. 4C). These data suggest that the lower number of mature OLs in P7 *Olig1-Ino80*-cKOs is a consequence of the lower number of PDGFRA^+^ OPs (Fig. 3C), not necessarily a change in the probability of individual OPs differentiating into OLs. By P14 the number-densities of CC1^+^ OLs in *Olig1-Ino80*-cKOs and littermate controls were no longer significantly different (Fig. 4E). Therefore, INO80 ablation leads to a diminished proliferation rate of OPs in the perinatal period, a reduced number of OPs and hence a reduced rate of accumulation of differentiated OLs – i.e. there is a developmental delay but ultimately no reduction in the number of differentiated OLs.

**Figure 4.**
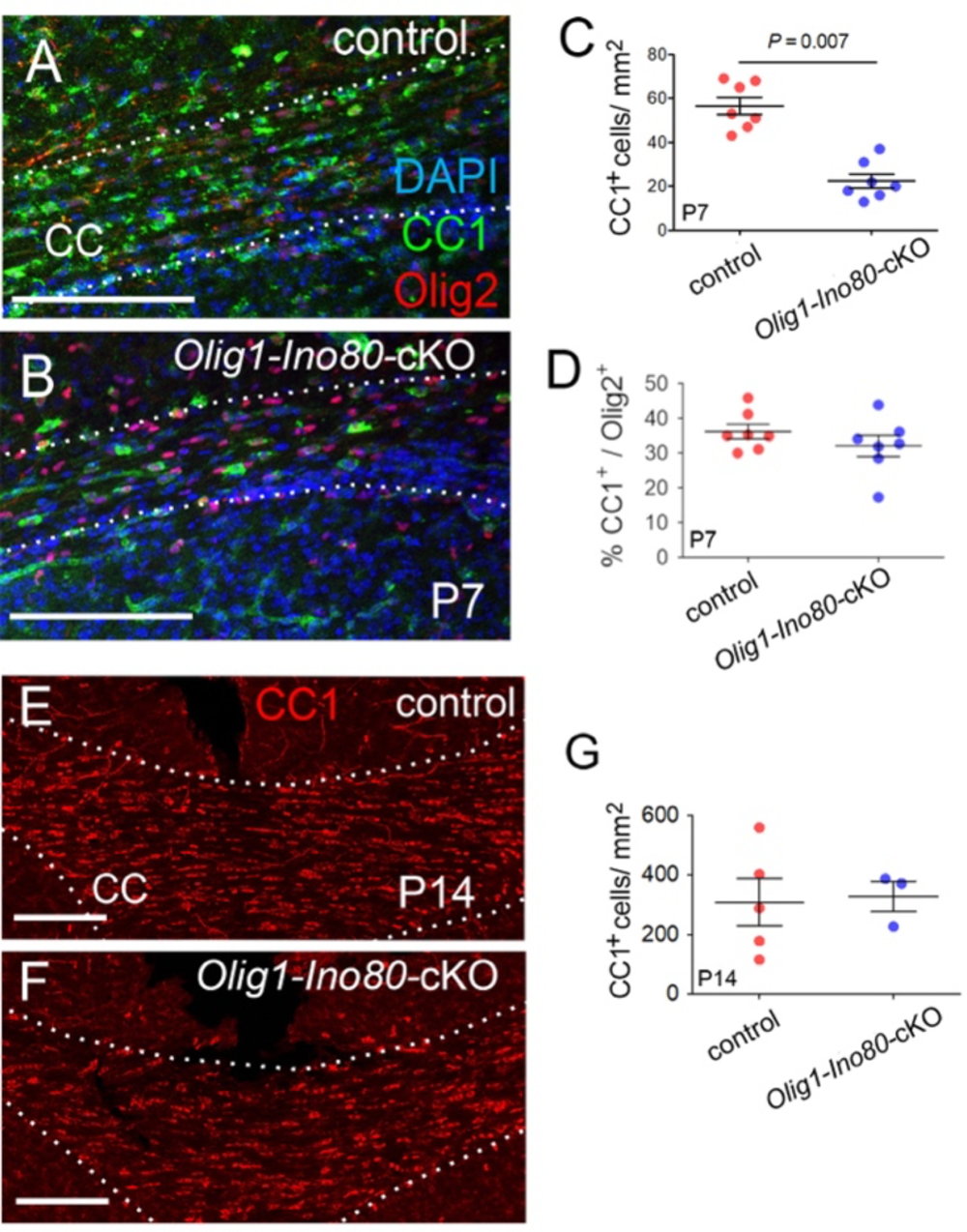
OL development in *Olig1-Ino80*-cKO mice. (**A,B**) P7 coronal sections of *Ino80^flx/flx^* control and *Olig1-Ino80*-cKO brains, immunolabelled for OLIG2 and the CC1 antigen and counterstained with DAPI. (**C**) Quantification of total (CC1+, OLIG2+) OLs and (**D**) (CC1+, OLIG2+) OLs normalized to OLIG2, to assess the fraction of mature OLs in the total OLIG2+ population. (**E,F**) CC1 immunolabelling of P14 coronal brain sections and (**G**) quantification of CC1+ OLs shows that there are similar numbers of differentiated CC1+ OLs in the corpus callosum of *Olig1-Ino80*-cKO and control littermates at P14. Data are presented as mean ± s.e.m. p-values according to Student’s t-test. Scale bars: 100 µm (**A, B**), 200 µm (**E, F**)

### INO80 ablation slows the cell cycle by prolonging G1

From the above it is clear that INO80 influences the duration of the OP cell cycle. To investigate this in more detail we turned to cumulative EdU labeling, which we have previously used to analyze the cell cycle of OPs in embryonic and postnatal mice (van Heyningen et al., 2001; Psachoulia et al., 2009; Young et al., 2013). We administered EdU to pregnant females (*Olig1-Cre: Ino80^flx/flx^* and *Ino80^flx/flx^* controls) in their drinking water, starting on embryonic day 17 (E17). We collected litters at various times thereafter so that different litters were exposed to EdU for different periods of time from 3 to 24 hours. At each time point we estimated the EdU labelling index of PDGFRA^+^ OPs (Fig. 5A,B). The labelling index increased linearly with time of exposure to EdU, as expected, for both *Olig1-Ino80*-cKO and control embryos; however, the rate of increase (i.e. slope of the line) was lower for *Olig1-Ino80*-cKOs than for controls (Fig. 5B), indicating a slower cell cycle. From these cumulative labeling data we can estimate both the duration of S-phase (T_S_) and the total cell cycle time (T_C_); T_S_ was 13.5 ± 2.0 hours in controls versus 16.7 ± 1.1 hours in *Ino80*-cKOs, while T_C_ was 76.6 ± 15 hours in controls versus 179 ± 17 hours in *Olig1-Ino80*-cKOs (Fig. 5E). This calculation (see Methods) assumes uniformly cycling populations of OPs in both controls and *Olig1-Ino80*-cKOs; this condition is not satisfied in *Olig1-Ino80*-cKOs, because we estimated the efficiency of CRE-mediated *Ino80* deletion to be ∼85% based on recombination of *Rosa-YFP,* or ∼75% based on genomic qPCR (Supplementary Fig. S1), so there will be at least two populations of OPs – recombined and unrecombined. Our estimate of T_C_ in *Olig1-Ino80*-cKOs is therefore likely to be an under-estimate. In any case we can conclude that most of the large increase in T_C_ occurs outside of S-phase, in G2/M/G1. To assess the duration of G2, we labelled dividing cells in newborn (P0) mice acutely with a single intra-peritoneal injection of EdU, sacrificed them one or four hours later and co-immunolabeled forebrain sections for OLIG2 and phospho-histone H3 (pH3), together with histochemistry for EdU (Fig. 5C,D). pH3 is a specific marker of M-phase nuclei, so OLIG2^+^ OPs that had been in S-phase at the time of EdU injection, but had not yet traversed G2 into M, were EdU^+^, pH3-negative, whereas OPs that had been through both S- and G2-phases and had entered M-phase were EdU^+^, pH3^+^. A one-hour chase following EdU labelling did not result in any OLIG2^+^, EdU^+^ OPs becoming pH3^+^ (i.e. G2 > 1 hour), whereas after the 4-hour chase a fraction of EdU^+^ OPs in both *Olig1-Ino80*-cKOs and controls were pH3^+^ (i.e. G2 < 4 hours in both cKOs and controls) (Fig. 5C). The fraction of EdU^+^ OPs that entered M-phase after the 4-hour chase was reduced in *Olig1-Ino80*-cKOs versus controls (6.0% ± 0.5% versus 16.5% ± 1.5%), indicating that the duration of G2 increases in the absence of INO80 (Fig. 5C). However, because of the relatively short duration of G2, even in the absence of INO80, it is clear that the large increase in T_C_ is due mainly to a greatly extended G1, from ∼60 hours in control mice to >158 hours in *Olig1-Ino80*-cKOs.

**Figure 5.**
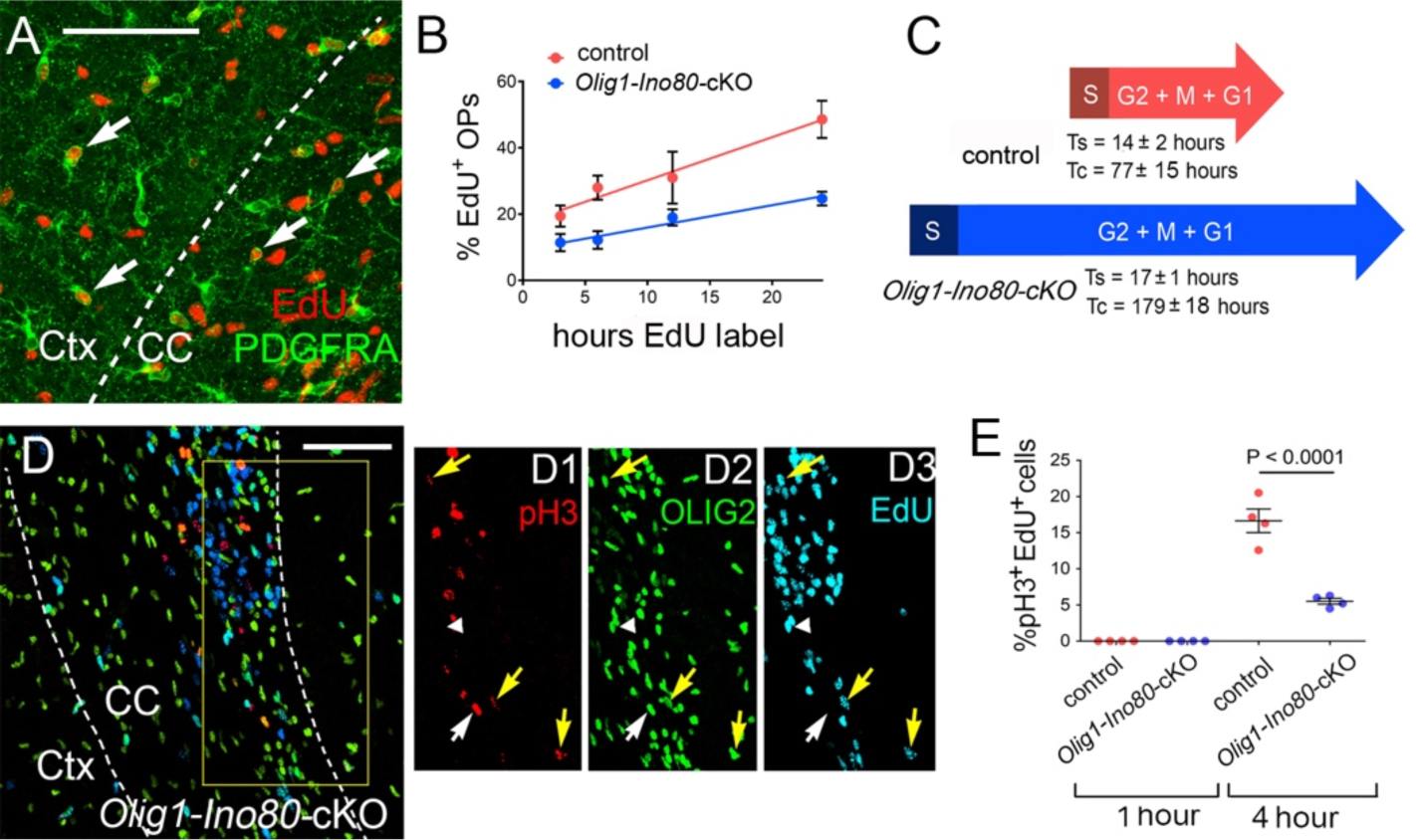
Loss of INO80 slows the OP cell cycle. We exposed pregnant female mice to EdU in their drinking water for various times, starting on embryonic day 17 (E17) until birth (∼E18) (cumulative EdU labelling). (**A**) Representative forebrain section of a neonatal (P0) *Olig1-Cre: Ino80^flx/flx^* (*Olig1-Ino80-cKO*) mouse imunolabelled for PDGFRA, along with histochemical detection of EdU. Dotted line indicates the boundary of neocortex. (Ctx) and corpus callosum (CC). Arrows indicate (EdU+, PDGFRA+) proliferative OPs. (**B**) We plotted the fraction of PDGFRA^+^ OPs that incorporated EdU in the CC as a function of EdU exposure time for *Olig1-Ino80-cKOs* and *Ino80^flx/flx^* controls. OPs in *Olig1-Ino80*-cKOs incorporated EdU at a lower rate than their littermate controls at all EdU exposure times. (**C**) From the slope and y-intercept we calculated the cell cycle times and durations of S-phase in *Olig1-Ino80*-cKOs and littermate controls (Methods). S-phase was little changed but the duration of G2+M+G1 was more than doubled from ∼77 h to >200 h in *Olig1-Ino80*-cKOs (see Methods). (**D**) To estimate the duration of G2-phase, we gave neonatal pups a single intra-peritoneal injection of EdU followed by a chase of 1 or 4 hours before immunolabeling forebrain sections with anti-OLIG2 and anti-phospho-H3 (pH3) together with EdU histochemistry. Separate channels for pH3 (red), OLIG2 (green) and EdU (blue) are shown in **D1**-**D3**. (**E**) Quantifying the fraction of OLIG2+ cells that was (pH3+, EdU+) showed that no cells co-labelled for pH3 and EdU after a one-hour chase, but that after a 4-hour chase a fraction of (OLIG2^+^, EdU^+^) OPs in the CC was also pH3^+^ and this fraction was greater in control mice than in *Olig1-Ino80*-cKOs, indicating that the duration of G2 was modestly increased in the absence of *Ino80* (estimated increase from ∼1.8 to ∼3 hours; see Methods). Therefore, most of the increase in cell cycle time T_C_ is because of a greatly extended G1-phase. Statistical significance is based on Student’s t-test. Scale bars: 50 µm (**A**); 100 µm (**D**).

### Transcriptomic analysis of INO80-deficient OPs indicates broad changes in OP gene expression

To gain a better insight into the transcriptional pathways and genes regulated by the INO80 complex, we conducted an RNAseq transcriptomic analysis using immunopanned OPs from P6-7 neonatal *Olig1-Ino80*-cKO and control mice. Dimensional reduction with principal component analysis (PCA) indicated cluster segregation based on genotype (Fig. 6A, Supplementary Fig. S2). The top differentially expressed genes highlighted a number of transcription factors that were down-regulated after *Ino80* gene deletion, including *Foxd1*, *Hoxa10*, *Pax3* and *Lef1* along with a number of cell adhesion genes including *Pcdha5*, *Pcdha9* and *Pcdha12* (Fig. 6A). Gene set enrichment analysis (GSEA) revealed that histone methylation genes were broadly down-regulated following *Ino80* gene deletion (Fig. 6B).

**Figure 6.**
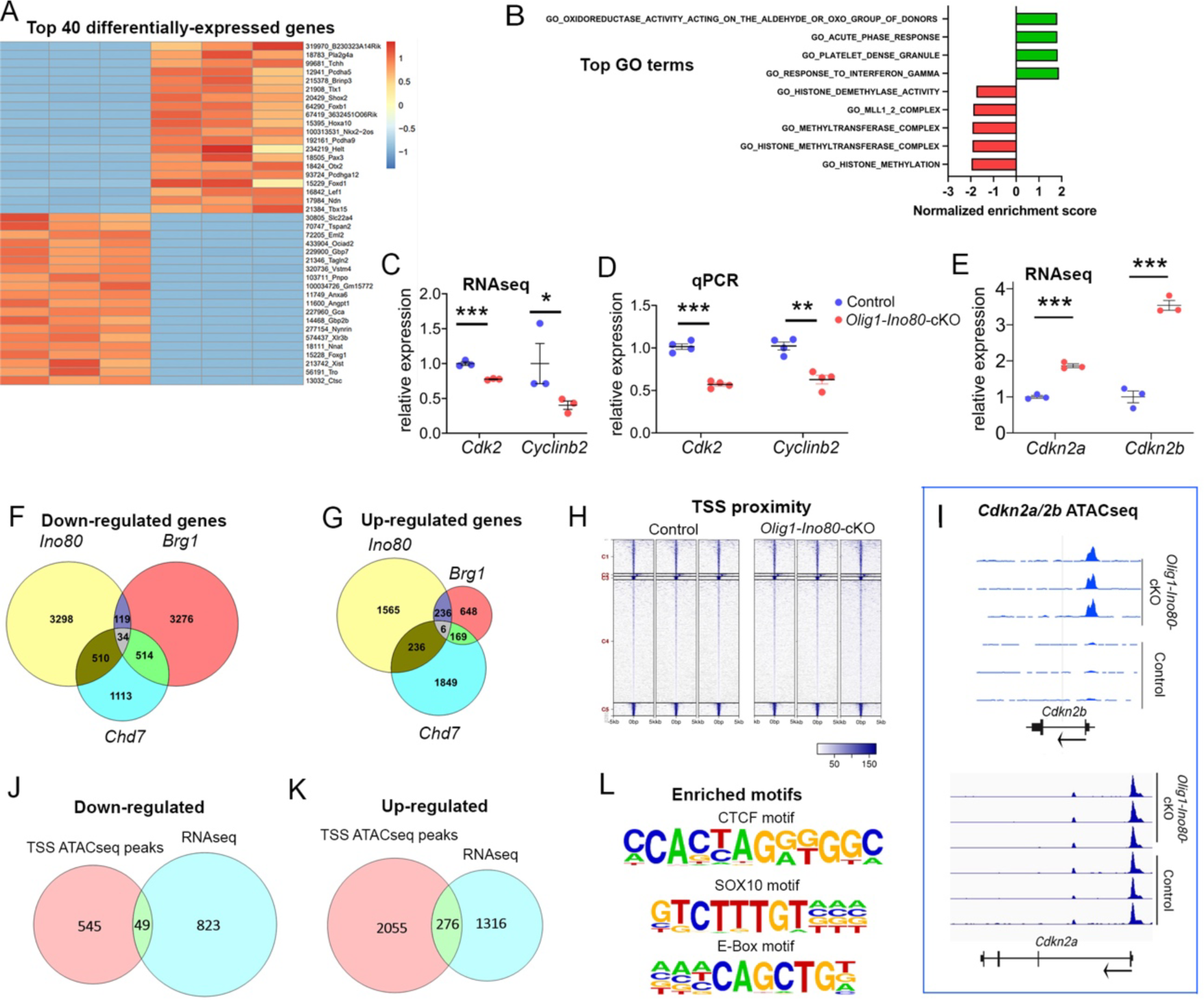
Transcriptomic and genome-wide chromatin accessibility changes in *Olig1-Ino80*-cKO OPs. OPs were immunopurified from P7 *Olig1-Ino80*-cKO and *Ino80^flx/flx^*control littermates (n = 3 per genotype), expanded in vitro and treated with 4-hydroxy-tamoxifen when the cultures were sufficiently dense. RNAseq analysis of these samples identified a number of differentially expressed genes (DEG). (**A**) The top 40 genes up- or down-regulated in *Olig1-Ino80*-cKO OPs are represented as a heat map [colour bars indicate log_2_(fold change)]. (**B**) Gene set enrichment analysis (GSEA) revealed significant down-regulation in *Olig1-Ino80*-cKO OPs of gene sets associated with histone methylation. (**C**) RNAseq analysis of the expression of known E2F target genes revealed significant down-regulation of the positive cell cycle regulators *Cdk2* and *Cyclinb2*, and this was confirmed by qPCR (**D**). RNAseq also revealed that the negative cell cycle regulators *Cdkn2a* and *Cdkn2b* were down-regulated in *Olig1-Ino80*-cKO OPs relative to controls. (**F**,**G**) Comparative DEG analysis looking at (**F**) down-regulated and (**G**) up-regulated transcripts in our own *Olig1-Ino80*-cKO RNAseq dataset compared to published *Brg1*-cKO and *Chd7*-cKO datasets. There is rather little overlap among the genesets affected by INO80, CHD7 or BRG1 complexes. ATACseq chromatin accessibility analysis was conducted on the same OP samples. (**H**) Global analysis of genome wide TSS chromatin accessibility revealed no overt differences in TSS occupancy in OPs of *Olig1-Ino80*-cKOs and littermate controls. (**I**) ATACseq peaks along the *Cdkn2b* gene reveal increased chromatin accessibility at the TSS in *Olig1-Ino80*-cKO compared to littermate controls, but no such effect along the *Cdkn2a* gene (arrows indicate direction of transcription). (**J**,**K**) There was no correlation between DEGs and TSS ATACseq peaks for the majority of DEGs with minimal overlap in (**J**) down-regulated DEGs and (**K**) up-regulated DEGs. (**L**) Motif analysis of significantly down-regulated ATAC-seq peaks revealed a significant enrichment of CTCF, SOX10 and E-Box motifs in the down-regulated fraction.

As GSEA did not flag cell cycle-associated genesets as significantly altered in our *Olig1-Ino80*-cKO OPs, we examined individual genes known to be involved in cell cycle control and G1 arrest. INO80 has been previously identified as an important regulator of S-phase entry via remodeling of histones at the transcription start sites of E2F target genes (Knezevic et al., 2018) – a well studied regulatory mechanism of the G1- to S-phase transition. Furthermore, *Ino80* deletion in other tissues has been shown to alter both E2F target gene expression and cell proliferation (Rhee et al., 2018). Our RNAseq dataset revealed that the E2F target genes *Cdk2* and *Cyclinb2* were down-regulated in *Ino80*-cKO OPs (Fig. 6C) which was confirmed by qPCR from independently isolated P7 OPs (Fig. 6D). Furthermore, *Cdkn2a* and *Cdkn2b*, coding for P14^ARF^/P16^INK4a^ and P15^INK4b^ cyclin kinase inhibitors respectively, were upregulated in *Olig1-Ino80*-cKO OPs (Fig. 6E). These are both E2F target genes involved in G1 arrest and senescence. This suggests that INO80-dependent slowing of the cell cycle is linked to mis-regulation of E2F target genes at the G1–> S transition checkpoint.

*Ino80* deletion did not appear to affect OP differentiation directly (see above) – unlike deletion of *Brg1* (Yu et al., 2013) or *Chd7* (He et al., 2016) – suggesting that the INO80 complex might influence a distinct set of genes. We therefore compared the published RNAseq datasets (Yu et al., 2013; He et al., 2016) with our own, to explore the functional overlap among different CRCs in OPs. This revealed rather little overlap in gene sets that were down- and up-regulated following loss of INO80, BRG1 or CHD7 (Fig. 6F,G). For example, of 4061 genes significantly up-regulated in either *Olig1-Ino80*-cKO or *Chd7*-cKO, only 242 (6%) were up-regulated in both. Of 2860 up-regulated in either *Olig1-Ino80*-cKO or *Brg1*-cKO, 242 (8%) were up-regulated in both. Of 3144 up-regulated in either *Brg1-cKO* or *Chd7*-cKO, 175 (6%) were up-regulated in both (Fig. 6G). Only 6 of 4709 genes (∼0.1%) were significantly up-regulated and 34 of 8864 genes (0.4%) were significantly down-regulated in all three cKOs (Fig. 6F,G). This suggests that the BAF, CHD7 and INO80 CRCs regulate largely separate transcriptional programmes in OPs.

### ATACseq analysis of Olig1-Ino80-cKO OPs indicates that chromatin accessibility does not account for the majority of transcriptional changes

We asked whether changes in chromatin accessibility were responsible for the transcriptional changes observed in *Olig1-Ino80*-cKO OPs. ATAC (assay for transposase-accessible chromatin) is a genome-wide method for tagging exposed regions of DNA, allowing a genomic map of nucleosome occupancy to be established by DNA sequencing (ATACseq). We immunopurified P6-P7 cortical OPs from *Olig1-Ino80*-cKO and control mice and compared them by ATACseq. This revealed no obvious systematic changes in nucleosome occupancy around transcription start sites (TSS) in *Olig1-Ino80*-cKO OPs relative to control OPs (Fig 6H). Of the differentially expressed E2F target genes that we identified, only *Cdkn2b* had significantly increased TSS chromatin accessibility in the *Ino80*-cKO OPs (Fig. 6I), consistent with its increased RNA abundance (Fig. 6E). We looked for a more general correspondence between TSS nucleosome occupancy and gene expression levels based on our RNAseq dataset but, for the majority of genes, altered TSS nucleosome occupancy did not correlate with either increased or decreased gene expression (Fig. 6J,K). This suggests that changes in transcription rates resulting from *Ino80* deletion do not necessarily result from altered nucleosome positioning around gene promoters or proximal enhancers.

To explore whether INO80 alters accessibility of DNA-binding sites for specific OL lineage transcription factors (TFs), we conducted a DNA sequence motif enrichment analysis on ATAC-seq peaks that we found to be reduced in *Olig1-Ino80*-cKO OPs relative to control OPs. This revealed an enrichment of a number of known transcription factor DNA-binding motifs including the CTCF zinc finger protein binding motif (enriched ∼11-fold; present in 10.13% of down-regulated ATAC peaks compared to 0.92% of all other peaks), the SOX10 motif (enriched 1.8-fold; present in 26.29% of down-regulated peaks compared to 14.53% of other peaks) and the E-box motif (enriched 1.9-fold; present in 30.15% of down-regulated peaks compared to 18.87% of other peaks); the E-box binds basic helix-loop-helix (bHLH) TFs including OLIG2 (Fig. 6L). These observations suggest that the INO80 complex might be targeted preferentially to promoters or enhancers that bind particular TFs, including certain OL lineage specific TFs, possibly because of physical interactions between specific TFs and the INO80 complex. A previous study demonstrated that OLIG2 interacts with BRG1, the BAF complex ATPase (Yu et al., 2013). To test for a direct physical interaction between OLIG2 and the INO80 complex we immunoprecipitated OLIG2 from cultured mouse OL lineage cells derived from E13.5 dissociated cortical precursor/stem cells and identified co-precipitated proteins by mass spectrometry. Of the 124 most abundant proteins in the precipitate, 14 were known CRC subunits, including INO80 and other components of the INO80 complex (Supplementary Fig. 3A-D). We looked for further evidence of an interaction between OLIG2 and the INO80 complex by proximity ligation assay (PLA) in cultured mouse OPs, using probe antibodies against INO80 and OLIG2 (Supplementary Fig. 3E). We obtained a positive signal in this PLA assay, indicating that INO80 and OLIG2 are co-located within 30-40 nm of each other in OPs and suggesting that OLIG2 associates physically with the INO80 complex, as with other CRCs.

### INO80 mediates nucleosomal histone H2A isoform exchange

It has been well documented that INO80 can facilitate exchange of nucleosome-bound histone H2A.Z for H2A (Papamichos-Chronakis et al., 2011; Alatwi and Downs, 2015; Brahma et al., 2017). Previous studies have shown that H2A.Z – H2A exchange at the TSS of E2F target genes is critical for proper transcription of these genes (Tarangelo et al., 2015; Knezevic et al., 2018). We therefore assessed the mobility of H2A.Z in cultured OPs to determine whether and how H2A.Z kinetics change in the presence or absence of INO80. We immunopurified and cultured OPs from *Ino80^flox/flox^* mice and expressed a H2A.Z-GFP fusion protein in these cells by transient co-transfection of *CMV-H2A.Z-GFP* together with either *CAG-Cre-T2A-mRuby* (*Ino80*-KO), or *CAG-mRuby* (control) (Fig. 7A-C). We identified transfected cells by mRuby intrinsic fluorescence and subjected them to fluorescent recovery after photobleaching (FRAP) or fluorescence loss in photobleaching (FLIP) assays to measure the intranuclear mobility of H2A.Z-GFP (Methods). The FRAP experiment involved inducing a single short-term photobleaching event within a discrete area of the cell nucleus and measuring the subsequent fluorescence recovery caused by movement of non-bleached H2A.Z-GFP into the bleached area over time (Fig. 7D,E). The calculated mobile fraction represents freely-diffusible H2A.Z-GFP and the immobile fraction represents DNA-bound H2A.Z-GFP. The FRAP experiment revealed that fluorescence recovery was more rapid in *Ino80*-KO nuclei than in control nuclei, indicating that the majority of H2A.Z-GFP was mobile within *Ino80*-KO nuclei, whereas the majority of H2A.Z-GFP in control nuclei was in the immobile DNA-bound fraction (Fig. 7E). The complementary FLIP experiment involved continuous illumination and photobleaching of a fixed area of the nucleus, while measuring loss of fluorescence intensity in a part of the nucleus outside the illuminated spot. This experiment demonstrated rapid nucleus-wide loss of H2A.Z-GFP fluorescence in *Ino80*-KO nuclei but not control nuclei, confirming that the majority of H2A.Z-GFP was freely diffusible in the absence of INO80 but DNA-bound in the presence of INO80 (Fig. 7F,G). Together, these experiments demonstrate that INO80 is critically important for proper H2A.Z incorporation into DNA-bound nucleosomes in OPs, as in other types of cell.

**Figure 7.**
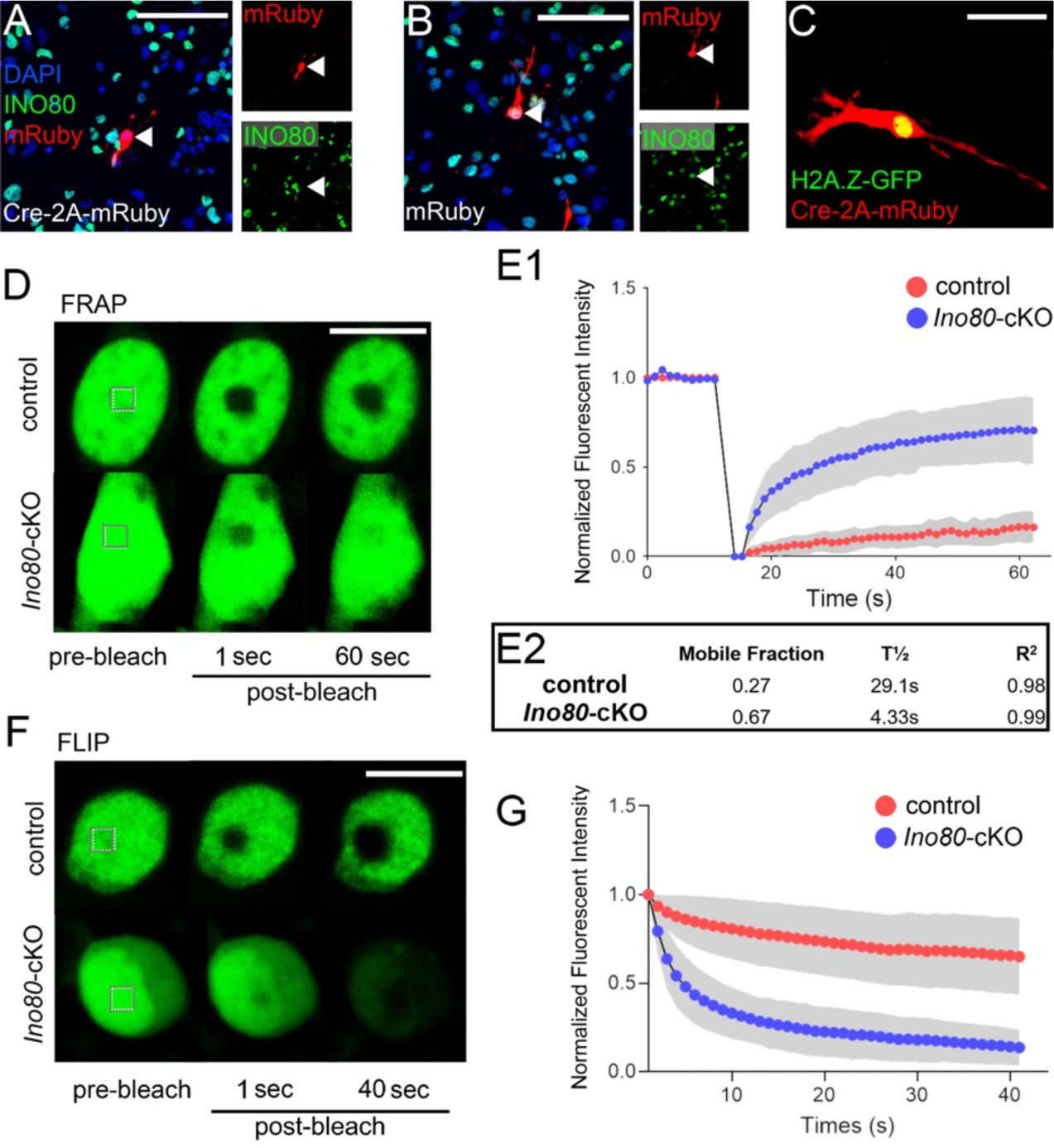
INO80-dependent mobility of histone H2A.Z in OP nuclei. (**A,B**) Immuno-purified OPs from *Ino80^flx/flx^* mice were transfected in culture with plasmids encoding H2A.Z-GFP fusion protein and either *Cre-T2A-mRuby* (**A**) or *mRuby* (**B**). 48 hours later the cells were fixed, immunolabelled for INO80 and counterstained with DAPI. Cells that expressed *Cre-T2A-mRuby* frequently did not immunolabel for INO80, confirming Cre-mediated recombination at the *Ino80* locus (*Ino80-cKO* cells, arrowhead in **A**). Cells that expressed mRuby alone invariably co-expressed INO80 (control cells, arrowhead in **B**). (**C**) Live cell fluorescence image of an OP co-expressing *H2A.Z-GFP* and *Cre-T2A-mRuby*, typical of those targeted for FRAP or FLIP analysis. (**D**) FRAP analysis of H2A.Z-GFP in a cultured OPs illustrates changes in H2A.Z dynamics over time in control and *Ino80*-cKO OPs. (**E1**) Time course of fluorescence intensity in the bleached area, showing more rapid fluorescence recovery in *Ino80*-cKO nuclei versus control nuclei. (**E2**) Calculated mobile H2A.Z fractions, half-times of recovery (T½) and R^2^ are tabulated. (**F**,**G**) FLIP analysis of H2A.Z-GFP in the nuclei of control and *Ino80*-cKO OPs, confirming higher mobility of histone H2A.Z in the absence of INO80. Scale bars: 50 µm (**A**, **B**), 20 µm (**C**), 5 µm (**D**, **F**).

## DISCUSSION

In this study we explored the role of the INO80-complex in the OL lineage. Using both CRISPR/CAS9-mediated *Ino80* gene disruption in cultured OPs and conditional *Ino80*-cKO in live mice, we uncovered a vital function of the INO80-complex in cell cycle progression of proliferating OPs, whereby absence of INO80 results in agreaterr extended G1 phase and a concomitant reduction in the rate of S-phase re-entry.

INO80 has been previously described as an important positive regulator of cell cycle in embryonic stem cells (Wang et al., 2014) and various cancers (Zhou et al., 2016b; Prendergast et al., 2020). We have shown, by a combination of cumulative and pulse-chase EdU labelling, Ki67 and phospho-histone H3 immunolabelling, that the cell cycle time T_C_ of perinatal OPs is substantially increased by loss of INO80 and that most of this increase reflects a much longer time spent in the G1-phase of the cycle (∼60 h in control mice vs ∼160 h in *Ino80*-cKOs). A previous study (Wang et al., 2014) showed that *Ino80* gene disruption in embryonic stem cells results in a protracted G1 phase, indicating that INO80 might be important for cell cycle progression across many proliferative cell types. The slowing of the OP cell cycle that we observed was more pronounced in the perinatal period than postnatally; at P14 the fraction of OPs that was G1-arrested (Ki67-negative) still tended to be less than littermate controls (p=0.066), although this did not reach the conventional threshold of significance (p≤0.05). Nevertheless, EdU labelling in young adult (P74-80) mice showed that OPs in *Pdgfra-Ino80*-cKOs had a lower EdU labelling index than controls (p=0.026), suggesting that *Ino80* deletion impacts the OP cell cycle at all ages.

We found that INO80 is dispensable for differentiation of perinatal OPs into CC1+/OLIG2+ OLs in vitro. Most other studies that have explored the function of CRCs in the OL lineage found inhibitory effects on OP differentiation into OLs. Deletion of the CRC ATPases CHD7, CHD8, EP400 and BRG1 all impacted OP differentiation (Yu et al., 2013; He et al., 2016; Elsesser et al., 2019), while CHD7 deletion was also shown to inhibit OP apoptosis through suppression of p53 expression (Marie et al., 2018). Therefore, the INO80 complex is the only one so far shown to impact the OP cell cycle specifically, without affecting differentiation directly. Collectively, these data point to a non-redundant role for each CRC in the OL lineage, possibly explained by the recruitment of each complex to a different set of gene targets and/or a specialized mechanism of action for each complex, involving distinct histone remodelling substrates or dynamics. The OL lineage, being one of the better-understood of all cell lineages, is an excellent model system in which to compare the influences of different CRCs, hence to understand their distinct functional roles and how these are integrated to control the dynamic transcriptional programmes that direct the development and physiology of all cells.

To try to explain the cell cycle phenotype observed in *Ino80*-cKO OPs, we examined the influence of the INO80-CRC on chromatin accessibility and gene transcription. INO80 has previously been reported to down-regulate the expression of E2F target genes (Rhee et al., 2018). In genome-wide analyses we did not detect any significant differential expression of gene sets identified as E2F targets, or associated broadly with cell proliferation or cell cycle dynamics. However, we did identify dys-regulation of a number of selected E2F target genes, including a significant increase in expression of *Cdkn2b*, which encodes p15^INK4b^, a CDK inhibitor that is well known to promote G1 arrest. Furthermore, ATAC-seq revealed a loss of the nucleosome free region around the *Cdkn2b* promoter, indicative of gene suppression. We looked genome-wide for a correlation between loss of ATAC-seq peaks at the TSS of genes and altered expression of the same genes in *Ino80*-cKO OPs, but could find no significant correlation, indicating that INO80-dependent differential gene expression was not due to reduced chromatin accessibility around the TSS. Furthermore, we did not observe any genome-wide differences between *Ino80*-cKOs and controls in nucleosome positioning within the bodies of genes, possibly because there is redundancy between the INO80 complex and other CRCs for this function. These observations indicate that INO80-dependent changes in nucleosome positioning around a gene’s proximal promoter is not critical for gene transcription and suggests that the transcriptional changes we observed following INO80 deletion are a result of a different mechanism or that INO80 acts through distal enhancers or other regulatory elements outside of the region we examined.

Motif analysis identified a number of transcription factor binding motifs that were enriched in genomic regions corresponding to ATAC-seq peaks that were reduced in *Ino80*-cKO samples compared to controls (i.e. chromatin that was less accessible in *Ino80*-cKO than control). One of these was the CTCF binding site. CTCF (CCCTC-binding factor) can occupy ∼60,000 sites across the mammalian genome, providing nucleosome positioning signals that help to define chromatin architecture and can activate or repress promoters and insulate enhancers (Krivega and Dean, 2017). Knock-out studies have demonstrated that CTCF is important for controlling chromatin accessibility, and interactions between CTCF and the BAF and ISWI CRCs have been demonstrated in previous studies (Marino et al., 2019; Wiechens et al., 2016). Our results indicate that, in OPs, DNA elements containing CTCF motifs are more likely to be accessible in the presence of INO80, relative to elements lacking CTCF motifs, so opening of chromatin by INO80 might depend partly on direct or indirect interactions between CTCF and INO80. SOX10-binding motifs and E-box motifs were also over-represented in chromatin regions rendered more accessible in the presence of INO80, adding to evidence that OL lineage-specific TFs can interact with a variety of CRCs (Yu et al., 2013; He et al., 2016; Elsesser et al., 2019). Supporting the ATAC-seq data, we found strong evidence from co-immunoprecipitation/ mass spectrometry that OLIG2 – an OL lineage-specific bHLH-TF that binds E-boxes – interacts with INO80 in OPs. It is possible that OL lineage progression relies on the interaction of OL lineage-specific TFs (such as SOX10 or OLIG2) with different CRCs at different stages of lineage progression, so as to influence expression of different sets of genes at different stages of OL development.

Beyond nucleosome positioning, the INO80-CRC has a well described role in H2A.Z to H2A histone exchange in nucleosomes (Brahma et al., 2017). In FRAP and FLIP experiments we identified a robust failure of global H2A.Z integration into DNA-bound histones following *Ino80* gene disruption, with 33% of H2A.Z-GFP bound to DNA compared to 73% in control OPs. This indicates that INO80 most likely influences gene expression in OPs through direct histone exchange rather than by influencing nucleosome positioning.

OPs divide rapidly during embryonic development in mice (van Heyningen et al., 2001) but their cell cycle slows down dramatically after birth, reaching a steady state in adulthood in which most OPs are arrested in early G1 but are still capable of re-entering the division cycle and generating myelinating OLs when required (Young et al., 2013). For example, we and others have shown that OP proliferation and OL production are stimulated and required during learning and/or memory formation in mice (McKenzie et al., 2014; Pan et al., 2020; Steadman et al., 2020; Shimizu et al., 2023), up to ∼90% of OPs can be stimulated to re-enter S-phase during training in an 8-arm radial maze task (Shimizu et al., 2023). Moreover, OPs are capable of re-entering a state of rapid division and differentiation to replace myelinating OLs that are lost in response to damage or disease. The intercellular signals that drive S-phase re-entry in these contexts are poorly understood, although PDGF is likely to play an important role (Calver et al., 1998; van Heyningen et al., 2001). Understanding how chromatin modifying proteins, including CRCs, drive OP division and differentiation in response to environmental signals is vital for building a picture of how these signals are integrated to regulate OL lineage dynamics in the healthy normal CNS and during disease.

## METHODS

### Mice

Animals were maintained on a 12 hour light/dark cycle with free access to food and water. Animal husbandry and procedures were pre-approved by the UCL Animal Welfare and Ethical Research Board (AWERB) and conformed to the the UK Animals (Scientific Procedures) Act 1986 and its amendment regulations 2012.

A “floxed” *Ino80* mouse line carrying *loxP* sites flanking exons 2-4 (*Ino80^flx^)* was obtained from Jackson Laboratories (stock no. 027920) (Qiu et al., 2016). This was crossed to an *Olig1-Cre* knock-in line (Lu et al., 2002) (obtained from David Rowitch, University of Cambridge, UK) to allow OL lineage-specific deletion of *Ino80* following tamoxifen administration. In addition, *Ino80^flx^* was crossed to *Pdgfra-CreER^T2^* transgenic mice (Rivers et al., 2008) to allow tamoxifen-dependent deletion of *Ino80* in OPs. In some experiments the *Rosa26-YFP* reporter strain (Srinivas et al., 2001) was introduced. Genotyping was by PCR (primer sequences in Table 1).

### EdU labelling in vivo

*Olig1-Cre: Ino80^flx/flx^* mice along with *Olig1-Cre: Ino80^flx/+^ or Ino80^flx/flx^* (no *Cre*) control littermates were collected at postnatal day 0 (P0), P7 and P14 after timed-mating. Animals were anesthetized with pentobarbital and intracardially perfused with PBS and 4% (w/v) paraformaldehyde (PFA). The perinatal mice received a single subcutaneous injection of 5-ethynyl-2′-deoxyuridine (EdU, 100 mg/kg body weight) 4 h prior to perfusion-fixation. For cumulative EdU labelling of perinatal embryos, pregnant females were injected intra-peritoneally once or multiple times at 2 h intervals with EdU (20 mg/kg), starting early on embryonic day 17 (E17) and continuing up to 24 h. Embryos were rapidly removed 2 h after the final injection and their brains removed and fixed in 4% PFA overnight at 4°C.

In the cumulative labelling method (Nowakowski et al., 1989; Psaschoulia et al., 2009), EdU labelling index (L.I., the fraction of PDFRA^+^ OPs that is also EdU^+^) is plotted on the y-axis as a function of EdU labelling time (T), giving a straight line of gradient *m* and y-intercept *y_0_*, until all cycling cells have incorporated EdU when the line reaches an abrupt plateau. The fraction of all OPs that is cycling (L.I. at plateau) is called the growth fraction (*G*). From these plots we can calculate cell cycle time T_C_ = *G*/*m* and S-phase duration T_S_ = *y_0_.G/m* (Nowakowski et al., 1989). We previously determined that *G* in normal perinatal mice is 100% and that T_C_ at E17 was ∼100 ± 17 h (van Heyningen et al., 2001), not very far off our present estimate of 77 ± 1 h (Fig. 5C, extrapolated from Fig.5B). We could not determine *G* in *Olig1-Ino80*-cKOs since we did not label sufficiently long with EdU (Fig. 5B). If we assume *G* = 100% then T_C_ is ∼179 h (Fig. 5C), but that is an average for the recombined (INO80-null) and unrecombined (wild type) OP sub-populations, the recombination efficiency in *Olig1-Ino80*-cKOs lies in the range 75%-85% (Supplementary Fig. S1C, D). Assuming the division rate of the unrecombined OP sub-population is the same as in CRE-negative control OPs, then our estimate of T_C_ in INO80-null OPs is an under-estimate. If we assume that 20% of OPs in *Olig1-Ino80*-cKOs are *not* recombined, hence dividing normally with T_C_ ∼77 h, then the recombined INO80-null sub-population would have to be dividing with T_C_ ∼[179 – (77×0.2)] / 0.8 = 205 h in order to give our measured *average* T_C_ of ∼179 h.

To analyze adult OPs, tamoxifen (300 mg/kg body weight in corn oil) was administered to *Pdgfra-CreER^T2^*: *Ino80^flx/flx^* mice by oral gavage on four consecutive days starting on P60 (Rivers et al., 2008) and EdU (0.2 mg/ml) was administered in their drinking water during the 7 days prior to perfusion on P80.

### Cell culture

Cells were cultured in basal medium consisting of Dulbecco’s Modified Eagle’s Medium (DMEM)/ F12 (ThermoFisher) supplemented with Glutamax (1x), N-2 supplement (0.5x), B-27 supplement (0.5x), non-essential amino acids (1x), sodium pyruvate (1x), penicillin/ streptomycin (1x), *N*-acetyl-*L*-cysteine (5 µg/ml) and recombinant human insulin (5 µg/ml). OPs were obtained by immuno-panning from single-cell suspensions of dissociated P6-7 mouse cortices using anti-PDGFRA immunoglobulin (558774, BD Pharmingen) as previously described (Emery and Dugas, 2013). Purified OPs were expanded as adherent monolayers on poly-D-lysine (PDL)-coated wells in basal medium with added PDGF-AA (10 ng/ml), FGF-2 (10 ng/ml) and NT3 (1 ng/ml). In some experiments OPs were cultured in basal medium containing tri-iodothyronine (T3) (40 ng/ml) to promote OL differentiation. Guide RNA (gRNA) sequences targeting INO80 were designed using benchling software (Table 1) and cloned into the PX459 *CRISPR/Cas9-2A-Puro* vector. pSpCas9(BB)-2A-Puro (PX459) V2.0 was a gift from Feng Zhang (Broad Institute, Boston, USA) (Addgene plasmid # 62988; http://n2t.net/addgene:62988; RRID:Addgene_62988). Transient transfection of PX459 into OPs was conducted using Lipofectamine 3000 (Thermo Fisher). Following transfection, productively transfected OPs were selected by supplementing the culture medium with 0.7 µg/ml puromycin for 3 days post-transfection. Cells were collected for DNA extraction to assess gRNA efficiency using the “surveyor” assay (Ran et al., 2013) with primers flanking the cut site (Table 1).

To delete *Ino80* from OL lineage cells in vitro, OPs were immuno-panned from P7 *Pdgfra-CreER^T2^*: *Ino80^flx/flx^* mice or *Ino80^flx/flx^* controls and treated with 2 µM of 4-hydroxy-tamoxifen in the culture medium for 3 days, refreshing the medium each day.

Neural stem cells (NSCs) were obtained from E13.5 mouse cortices and maintained in basal medium containing EGF (10 ng/ml) and FGF2 (10 ng/ml). OL differentiation was induced by replacing EGF and FGF2 in the medium with tri-iodothyronine (T3) (40 ng/ml) for 5 days.

### Western blots

Cells were lysed on ice in lysis buffer - 50 mM Tris-HCl (pH 7.5), 150 mM NaCl, 1% (v/v) NP-40, 0.5% (w/v) Na deoxycholate and 1x protease inhibitor mix (Roche). Cell debis was pelleted by centrifugation (30,000 x g) and the supernatant decanted for Western blot experiments. Protein concentration was quantified using the Bradford protein assay. 20 µg of protein was boiled in 4x laemmli sample buffer (Bio-Rad) for 5 mins and subject to SDS-PAGE and transferred onto a 0.45 µm PVDF membrane (Amersham) overnight (30V) using high molecular weight transfer buffer – 39 mM glycine, 48 mM Tris base, 10% methanol, 0.1% SDS. Membranes were subsequently blocked in 5% (w/v) skim milk PBST and incubated overnight in primary antibodies (Table 1), probed with an appropriate secondary antibody and incubated in Clarity ECL (Bio-Rad) prior to exposure to photographic film.

### Immunohistochemistry and cell counts

For *in vitro* experiments, cells were cultured on glass coverslips, washed in PBS containing Ca^2+^/Mg^2+^ and fixed in 4% (w/v) paraformaldehyde (PFA) for 10 mins. For in vivo mouse experiments, postnatal mice were perfused with cold PBS and 4% (w/v) PFA as previously described (Rivers et al., 2008). Brains were dissected and post-fixed for 2 h prior to incubation in 20% (w/v) sucrose cryoprotectant. Mouse brain tissue was embedded in OCT, frozen in an isopentane bath on dry ice and coronal cryosections (15 µm) collected on Superfrost® plus slides (Thermo Fisher). Fixed cultured cells and tissue sections were permeabilized and blocked in 0.1% (v/v) Triton X-100, 5% bovine serum in PBS for 1 h at 20-25°C, incubated overnight in primary antibodies (Table 1) diluted in 2% (v/v) bovine serum in PBS overnight at 4°C, then incubated in appropriate secondaries diluted in 2% bovine serum in PBS for 1 h at 20-25°C and counterstained with Hoechst 33257 (1:1000; Sigma). For brain tissue incubated in anti-INO80 antibody (ab118787; Abcam), prior to blocking, sections were subjected to an antigen retrieval step to expose the epitope. Sections were incubated for 5 min at 37°C in Proteinase K solution – 0.6 U/ml Proteinase K, 2.5% (v/v) glycerol, TE buffer (pH 8.0). Sections were immediately washed in 5% (v/v) bovine serum in PBS to inhibit the reaction. EdU staining was conducted using the EdU Click-iT assay (Thermofisher). Proximity ligation assays were performed on fixed cultured OPs using the Duolink® PLA kit (Sigma-Aldrich). This was conducted using anti-INO80 and anti-OLIG2 antibodies along with anti-SOX10 (positive control) and anti-PDGFRA (negative control) (antibody details in Table 1). Brain sections and cultured cells were counterstained with 300 µM of 4′,6-diamidino-2-phenylindole (DAPI) in PBS for 10 mins to label cell nuclei.

Immuno-labelled and EdU-labelled cells were counted in three 25 μm-thick coronal brain sections +1.7 mm to −0.3 mm relative to bregma in neonatal animals or +1.1 mm to −0.85 mm bregma in adults. Micrographs were taken using a 20x/ 0.50 N.A. plan-neofluar objective on a Zeiss LSM 880 Axio Imager 2 confocal microscope. For neonatal time-points, tiled images of the entire medial and lateral subcortical white matter including the corpus callosum were captured and for adult time-points, the corpus callosum between the medial limits of the lateral ventricles. Cells were counted with the aid of ImageJ image analysis software; the corpus callosum was delineated using anatomical references and the area measured. Cells were counted within this area and recorded as number-density (cells/ mm^2^).

### Rapid immunoprecipitation mass spectrometry of endogenous protein (RIMES)

RIMES cell processing, protein isolation and immunoprecipitation was as previously described (Mohammed et al., 2016) using 1×10^7^ OL-differentiated NSCs incubated with rabbit anti-OLIG2 (AB9610, Millipore) and using rabbit pre-immune IgG as control. Three replicates per group were prepared for proteomic analysis by mass spectrometry. Immunoprecipated proteins on beads were digested to peptides using trypsin. Peptides were desalted using C18+carbon top tips (TT2MC18.96, Glygen Corporation) and eluted with 70% (v/v) acetonitrile with 0.1% (v/v) formic acid.

Dried peptides were dissolved in 0.1% (v/v) trichloroacetic acid (TCA) and analyzed by low-flow liquid chromatography on an UltiMate^TM^ 3000 RSLCnano instrument, coupled on-line to a Q Exactive^TM^ Plus mass spectrometer (both Thermo Scientific). Gradient elution was from 3% to 35% (v/v) buffer B for 120 min at a flow rate 250 nL/min with buffer A being used to balance the mobile phase (buffer A was 0.1% (v/v) formic acid in water and B was 0.1% formic acid in ACN). The mass spectrometer was controlled by Xcalibur software (version 4.0) and operated in positive mode. The spray voltage was 1.95 kV and the capillary temperature was set to 255°C. The Q-Exactive Plus was operated in data-dependent mode with one survey MS scan followed by 15 MS/MS scans. The full scans were acquired in the mass analyzer at 375-1500 m/z with resolution of 70,000, and the MS/MS scans were obtained with a resolution of 17,500. The raw data files are available via the PRIDE database accession number XXXXXX.

MS raw files were converted into Mascot Generic Format using Mascot Distiller (version 2.5.1) and searched against the SwissProt database (December 2015 release) restricted to mouse entries using the Mascot search daemon (version 2.5.0) with a FDR of ∼1%. Allowed mass windows were 10 ppm and 25 mmu for parent and fragment mass-to-charge values, respectively. Variable modifications included in searches were oxidation of methionine, pyro-glu (N-term) and phosphorylation of serine, threonine and tyrosine. The mascot result (DAT) files were extracted into Excel files for further normalization and statistical analyses.

### FRAP and FLIP assays

OPs were immuno-purified from P6-P7 *Ino80^flx/flx^* mice, cultured on 40 mm coverslips and co-transfected with *pIN H2A.Z-GFP* (Addgene #15770) and *pCAG-Cre-T2A-mRuby2* (Addgene #102989). Control cultures were co-transfected with *pIN H2A.Z-GFP* and *pCAG-mRuby* lacking the *Cre-T2A* cassette. 48 hours post-transfection, fluorescence recovery after photo-bleaching (FRAP) or fluorescence loss in photobleaching (FLIP) was performed on a Zeiss LSM 880 scanning confocal microscope using the 488 nm line of the argon laser. Only cells expressing both mRuby2 and H2A.Z-GFP were analyzed. FLIP and FRAP were conducted with a Zeiss LSM780 confocal microscope using an EC Plan-Neofluar 40x/1.3 oil objective. <u>FRAP</u>: 10 pre-bleach images were obtained, then a single bleaching pulse was delivered at 100% laser power in a 2 µm^2^ circular area of the nucleus. Subsequent post-bleach imaging was conducted at 2 Hz for 62 sec. Fluorescent intensity of bleached, unbleached and non-nuclear background regions of interest (ROIs) were obtained using the Zen software package (Zeiss). <u>FLIP</u>: A continuous bleaching pulse was delivered at 100% laser power in a 2 µm^2^ area of the nucleus. Bleaching and imaging was performed at 2 Hz for 42 sec and fluorescent intensities of chronically bleached, unbleached and non-nuclear background ROIs were obtained using Zen software. FRAP and FLIP experiments were performed in triplicate and analyzed using EasyFRAP (Koulouras et al., 2018) to normalize data and extract mean fluorescence half-life T_1/2_ and mean mobile fraction of nuclear H2A.Z-GFP.

### Quantitative PCR

Quantitative PCR (qPCR) was conducted on OP cultures used for RNAseq experiments and YFP+ OPs isolated from postnatal day (P)7 *Olig1-Cre: Ino80^flx/flx^: Rosa26-YFP* and *Olig1-Cre: Ino80^flx/+^: Rosa26-YFP* mice. Tissue was lysed and homogenized with Trizol reagent (Invitrogen) and total RNA purified and used for cDNA synthesis using the superscript™ synthesis system (Thermo Fisher). Oligonucleotide qPCR primers are shown in Table 1. qPCR values were calculated using the relative standard curve method and samples normalized to *Gapdh*. At least three biological replicates for each genotype were analyzed.

### RNAseq and ATACseq

OPs were obtained from P6-7 *Olig1-CreER^T2^*: *Ino80^flx/flx^* mice or *Ino80^flx/flx^* controls and expanded in culture to obtain 2 x 10^6^ cells for bulk RNAseq analysis and 1 x 10^5^ cells for ATACseq analysis. 4-hydroxytamoxifen was added to the culture medium at a concentration of 2 µM. Cells for RNAseq were pelleted and snap-frozen; cells for ATACseq were harvested and prepared using the Active Motif proprietary method. 3 biological replicates from experimental and control groups were prepared in this fashion. Subsequent downstream sample preparation and analysis was conducted by Active Motif. <u>RNAseq:</u> RNA was analyzed using a BioAnalyzer and samples with 8.7-10.0 RNA integrity number values were used for library preparation using the Illumina TruSeq RNA Sample Preparation v2 Guide (Illumina). Libraries were sequenced on an Illumina NextSeq 500 as 42bp paired-end reads (PE42). Reads were aligned to the mouse genome (mm10) using the STAR algorithm with default settings and fragments overlapping genes were counted. Only paired reads both ends aligned were counted and gene with the same Entrez gene identifier were merged into a single gene. Differential analysis of gene fragment counts between experimental (*Olig1-CreER^T2^: Ino80^flx/flx^*) and control (*Ino80^flx/flx^*) groups was conducted using DESeq2. Gene Set Enrichment analysis (GSEA) was conducted using the DESeq2 normalized gene counts with the MSigDB’s C5 GO gene set collection. To compare our *Olig1-Ino80*-cKO RNAseq differentially regulated genes with *Brg1* cKO RNAseq (GSE42443) and *Chd7* cKO RNAseq (GSE72726) datasets, we obtained common down-regulated, up-regulated and uniquely differentially-regulated genes via DESeq2 analysis and represented the data in a Venn diagram. <u>ATACseq:</u> The ATACseq library was prepared using the Nextera DNA Library Prep Kit (Illumina) with the Active Motif proprietary method and sequenced on Illumina NextSeq 500 as 42bp paired-end reads (PE42). Reads were mapped to the mouse genome (mm10) using the BWA algorithm. Peak calling was conducted using the MACS2 algorithm. Fragment density analysis, data normalization and merged region analysis for peak metrics comparison between our 2 experimental groups was conducted using Active Motif proprietary software. Peaks were visualized using Bigwig output files with the Integrated Genome Browser (IGB). Motif analysis of statistically significant differential ATACseq signals was conducted using HOMER-based motif analysis.

## ACKNOWLEDGMENTS

We thank our colleagues at UCL and QMUL for help, advice and encouragement. We especially thank Matthew Grist for technical and administrative assistance. This work was supported by the UK Biotechnology and Biological Sciences Research Council (BB/S008934/1 to W.D.R. and H.L.) the Wellcome Trust (214286/Z/18/Z to W.D.R.) and a Sanming Project (SZSM201911003) funded by the Municipal Government of Shenzhen, China to W.D.R, H.L. and others.

## AUTHOR CONTRIBUTIONS

J.W., H.L. and W.D.R. conceived the project. W.D.R. and H.L. obtained funding and supervised the work. J.W. performed most of the experiments; Y.J. helped with the experiments involving OP immunopanning and cell culture. S.N. helped maintain mice. J.W., H.L. and W.D.R. interpreted the data. J.W. and W.D.R. wrote the paper with input from all authors. V.R. and P.C. performed mass spectrometry and data analysis at QMUL.

## Conflict of interest

The Authors declare no competing interests.

---

**Supplementary Figure 1.**
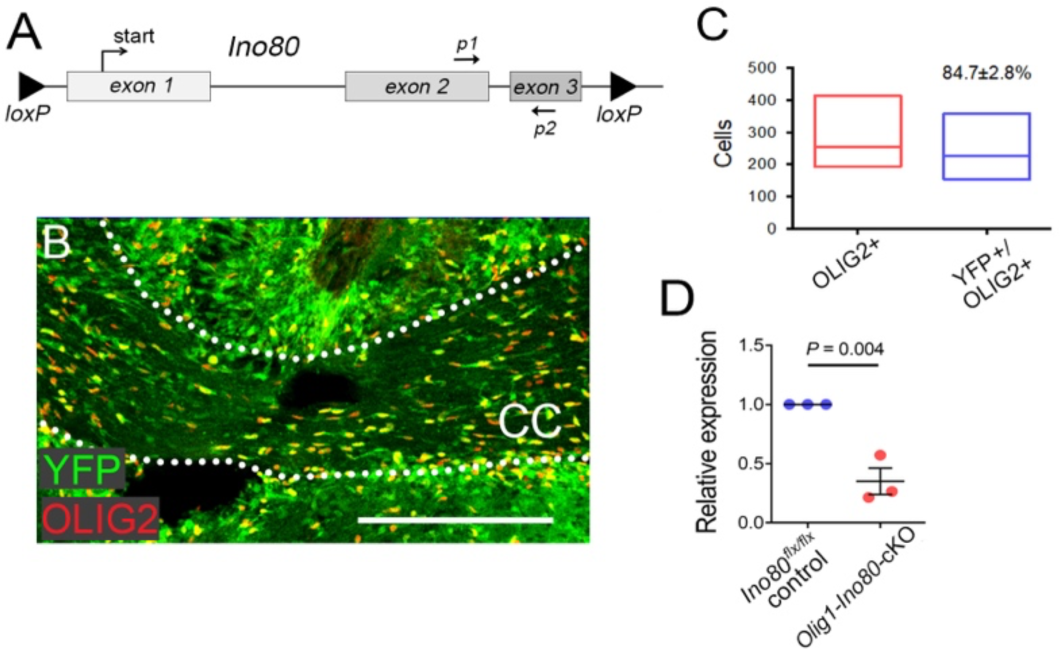
Recombination efficiency in *Ino80*-cKO mice. (**A**) Positions of *loxP* sites at the modified *Ino80^flx^*loci in transgenic mice. Also shown are the relative positions of PCR primers (P1, P2) used to assess Cre-mediated recombination in reverse transcription PCR (rtPCR) assays of purified mRNA. (**B**) Fluorescence image of the corpus callosum (CC, dotted lines) from a P7 *Olig1-Cre: Rosa26-YFP* mouse immunolabelled for GFP and OLIG2; double-labelling for OLIG2 and YFP reflects Cre-mediated recombination of the *Rosa26-YFP* reporter transgene in OL lineage cells. (**C**) Quantification of OLIG2^+^ and YFP^+^/OLIG2^+^ cells in P7 *Olig1-Cre: Rosa26-YFP* mice; recombination efficiency is ∼85% based on YFP expression. (**D**) Quantitative RT-PCR of RNA from OPs immuno-purified from P7 *Olig1-Ino80*-cKO and *Ino80^flx/flx^* control brains, using primers (*p1, p2*) positioned as in (**A**) suggests that CRE recombination efficiency is ∼75%. Error bars represent mean ± s.e.m. Statistical significance based on Student’s t-test. Scale bar: 100 µm (**B**).

**Supplementary Figure 2.**
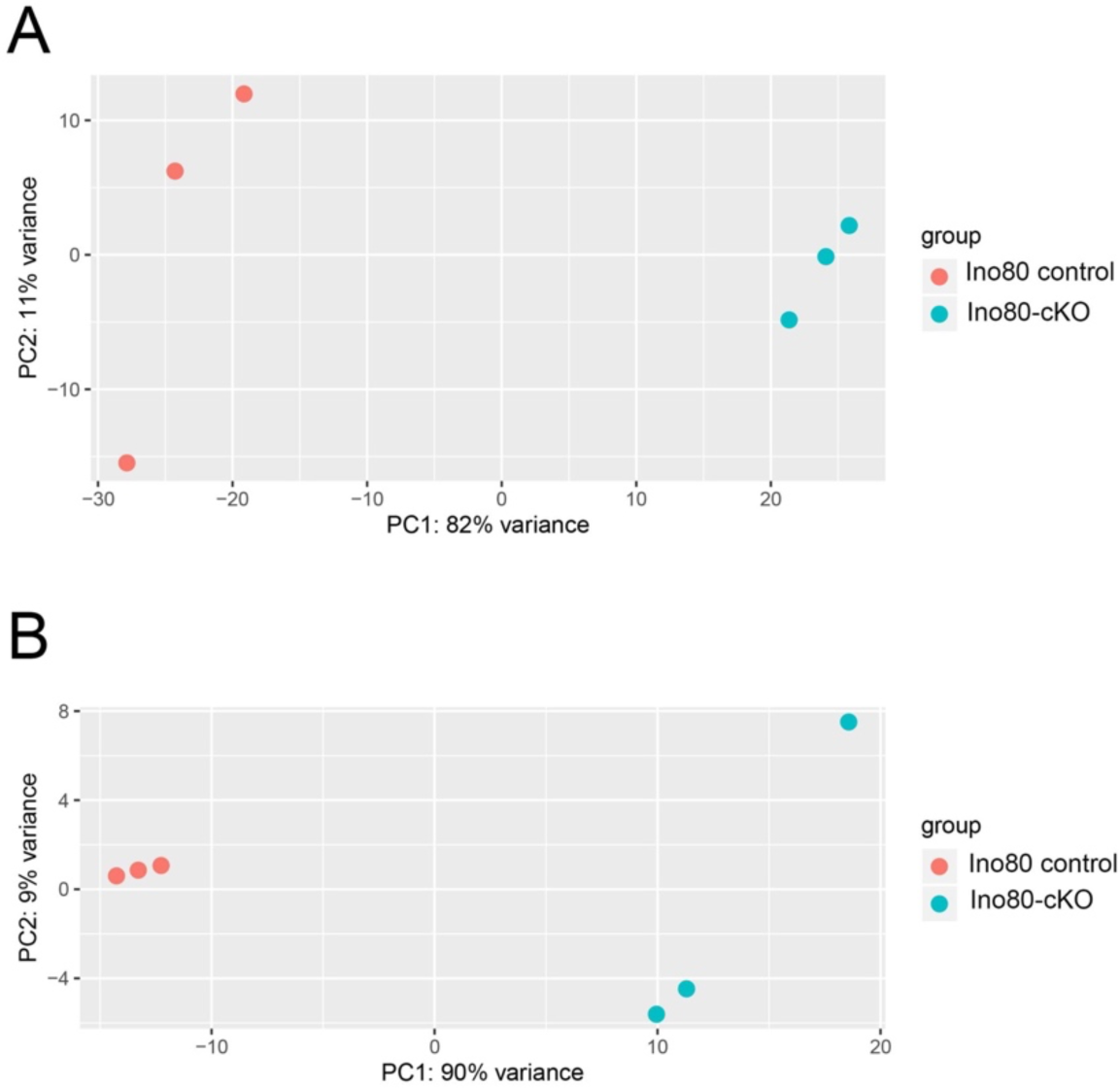
PCA plots of RNAseq and ATACseq datasets. PCA analysis of (**A**) RNAseq datasets and (**B**) ATACseq datasets from *Ino80*-cKO (green) and *Ino80* control (red) OP replicates.

**Supplementary Figure 3.**
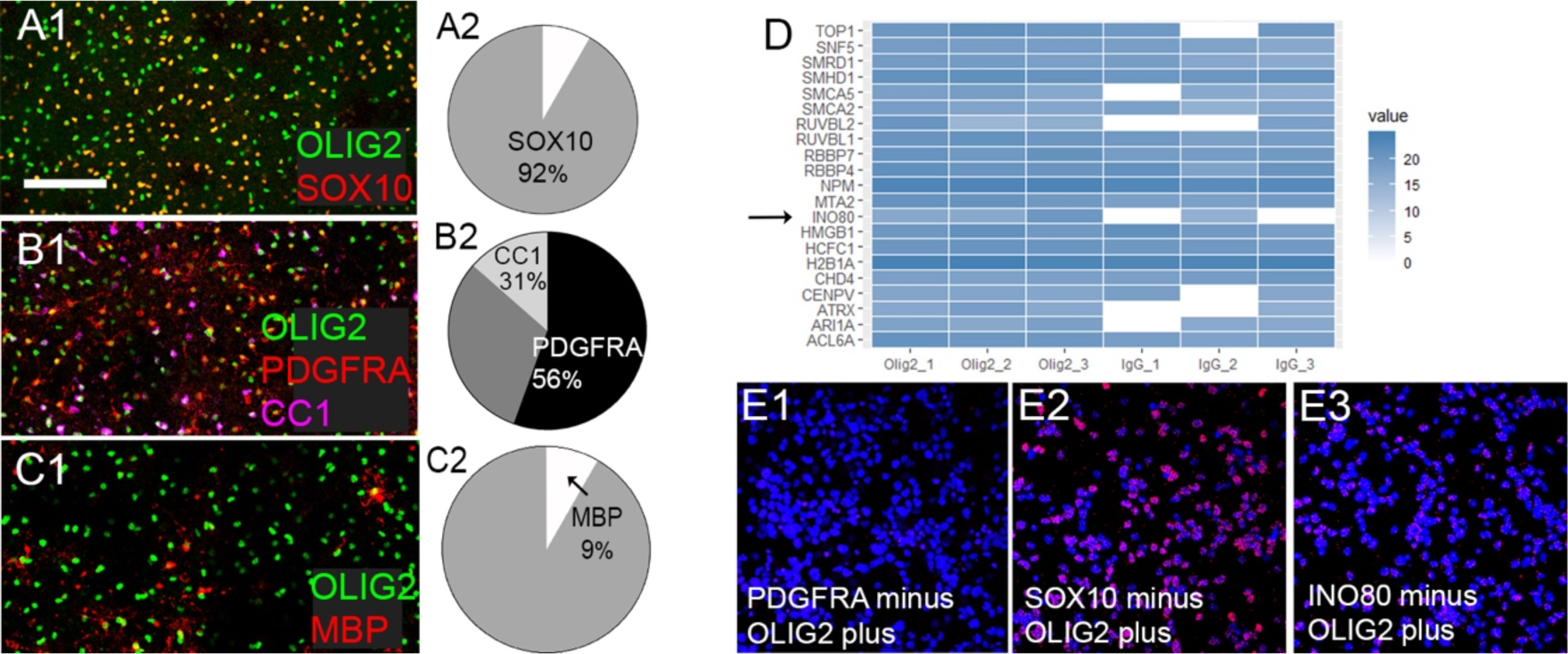
Protein-protein interactions between OLIG2 and components of chromatin remodeling complexes. Monolayer cultures were established from dissociated mouse E13.5 cortical hemispheres (neural stem cells, NSCs) and cultured in medium containing thyroid hormone (TH) to induce OL differentiation for 5 days before immuno-labelling for (**A1**) OLIG2 and Sox10, (**B1**) OLIG2, PDGFRA and CC1 antigen and (**C1**) OLIG2 and MBP. Pie charts illustrate the proportions of OLIG2^+^ cells that co-labelled with Sox10 (**A2**), Pdgfra and CC1 (**B2**) and MBP (**C2**). (**D**) Mass spectrometry was carried out on immuno-precipitates (IPs) from 5-day differentiated NSC-derived cultures. Around 175 proteins were identified that were >2-fold more abundant in all three replicate anti-OLIG2 IPs compared to negative control IgG Ips performed under identical conditions. Of these, 22 (∼12%) were known components of chromatin remodelling complexes (CRCs), including INO80 (arrow). (**E**) Proximity ligation assays provide further evidence of a close physical interaction between INO80 and OLIG2 in cultured OPs. Pdgfra-minus + OLIG2-plus probes provided a negative control (**E1**) and Sox10-minus + OLIG2-plus probes a positive control (**E2**). Red puncta indicate sites of close proximity indicating physical interactions between OLIG2 and Sox10 (**E2**), and between OLIG2 and INO80 (**E3**). Scale bar: 100 µm.

## Notes

### Competing Interest Statement

The authors have declared no competing interest.

